# *Tbx2a* modulates switching of opsin gene expression

**DOI:** 10.1101/676478

**Authors:** Benjamin A Sandkam, Laura Campello, Conor O’Brien, Sri Pratima Nandamuri, William Gammerdinger, Matthew Conte, Anand Swaroop, Karen L Carleton

## Abstract

Differences in sensory tuning are reported to maintain species boundaries and may even lead to speciation. Variation in the tuning of color vision is likely due to differences in the expression of opsin genes. Over 1,000 species of African cichlid fishes provide an excellent model system for studying the genetic basis of opsin gene expression because of the presence of seven distinct genes, yet individual species typically express only a divergent set of three opsins. The evolution of such shifts is expected to arise through either (a) two simultaneous regulatory changes (one for each opsin), or (b) one regulatory change that simultaneously promotes expression of one opsin while repressing another. Here, we used QTL analyses, genome sequencing, and gene expression studies to identify the transcription factor Tbx2a as likely driving a switch between *LWS* and *RH2* opsin expression. Binding sites for Tbx2a in the *LWS* promoter and the highly conserved Locus Control Region of *RH2* act to concurrently promote *LWS* expression while repressing *RH2* expression. Our data support the hypothesis that instant changes in visual tuning can be achieved by switching the expression of multiple genes by a single mutation and do not require independent changes in the regulatory regions of each opsin.

## INTRODUCTION

Color vision enhances the ability of organisms to detect signals from food, conspecifics, predators, and the environment, and can therefore assert a profound influence on fitness (1). Changes to the spectral tuning of visual signals are predicted to have dramatic effects on the evolutionary trajectory of species and even lead to speciation (2). It is thus of considerable interest to understand the genetic basis of evolutionary changes in visual sensitivity. The most variable component in tuning color vision are G-coupled protein receptors, cone opsins (3), which are expressed in the cone cells of the retina and, in most cases, are bound to the chromophore 11-*cis* retinal to form a specific visual pigment. The wavelength sensitivity of the visual pigment depends primarily upon the sequence of the opsin protein (4). Evolutionary changes in visual sensitivity were thought to be the result of changes in opsin amino acid sequence (4-6). However, accumulating evidence has shown that visual systems are frequently tuned more by changes in the relative expression of the opsin genes which result in larger shifts in tuning (7-11).

Modulation of visual sensitivity through shifts in expression requires the upregulation of one opsin coupled with concomitant downregulation of another (12-14). If each opsin is independently controlled by one or more transcription factors, simultaneous evolutionary changes in the expression of two opsins would be extremely unlikely. Differences in visual tuning among species would then require multiple substitutions that might be constrained by mutational dynamics. However, shifts in opsin expression could occur more quickly if a single regulatory element could down-regulate expression of one opsin while simultaneously up-regulating expression of another. Sharing regulatory elements across opsins would also facilitate correlated developmental shifts in opsin expression.

To determine whether such a common regulatory framework exists, we have examined the mechanisms of opsin gene expression in the radiation of cichlid fishes in East Africa. More than 1,000 closely-related species of African cichlid fishes represent an excellent model system for studying the genetic basis of opsin gene expression. The genome of each cichlid species encodes seven distinct opsin genes, yet an individual species typically expresses only three genes as adults, with the particular set of expressed genes being different among species (reviewed in 15). In addition, gene expression has been demonstrated to shift through development with species differences being the result of heterochronic changes to this developmental program (12, 16).

We previously analyzed a cross between *Aulonocara baenschi*, a Lake Malawi species with high expression of *RH2A* opsin and no expression of *LWS* opsin, and *Tramitichromis intermedius*, a species with low expression of *RH2A* but high expression of *LWS* (17). Among the *F*2 from the cross, we detected a strong negative correlation between expression of *LWS* and *RH2A* (*R*^2^ = −0.78) (18). We now report the identification of a *trans*-acting expression quantitative trait locus (eQTL) on LG10 that explains 48.6% of the variation in *LWS* expression and 57.1% of the variation in *RH2A*. The negative correlation and overlapping eQTLs suggest a common regulatory element that can simultaneously promote expression of *LWS* while repressing expression of *RH2A*. Our combined fine mapping, transcriptomics, and functional analyses show that the transcription factor Tbx2a likely controls the switch between *LWS* and *RH2A* expression.

## RESULTS

### The transcription factor Tbx2a is strongly correlated with the switch between LWS and RH2A opsin expression

In order to identify the factor responsible for switching *LWS*/*RH2A* expression between *A. baenschi* and *T. intermedius*, we first used microsatellite markers and previously developed RAD-seq markers (17) to fine map the *LWS*/*RH2A* eQTL to a region that spans 1.15 Mb and contains 31 genes (Fig. 1A & B, Supplementary Table 1). We predicted that the expression of candidate transcription factors which potentially regulate *LWS* should be correlated with *LWS* expression. In retinal transcriptomes from *T. intermedius* (N=2) and *A. baenschi* (N=4) (an average of 183 million reads) only three genes were differentially expressed between *A. baenschi* and *T. intermedius* with FPKM ratios greater than 2 (*adora2b* (5.04), *gjb1*, (2.13) and *Tbx2*a (105.95)) (Supplementary Table 1).

**Fig. 1.**
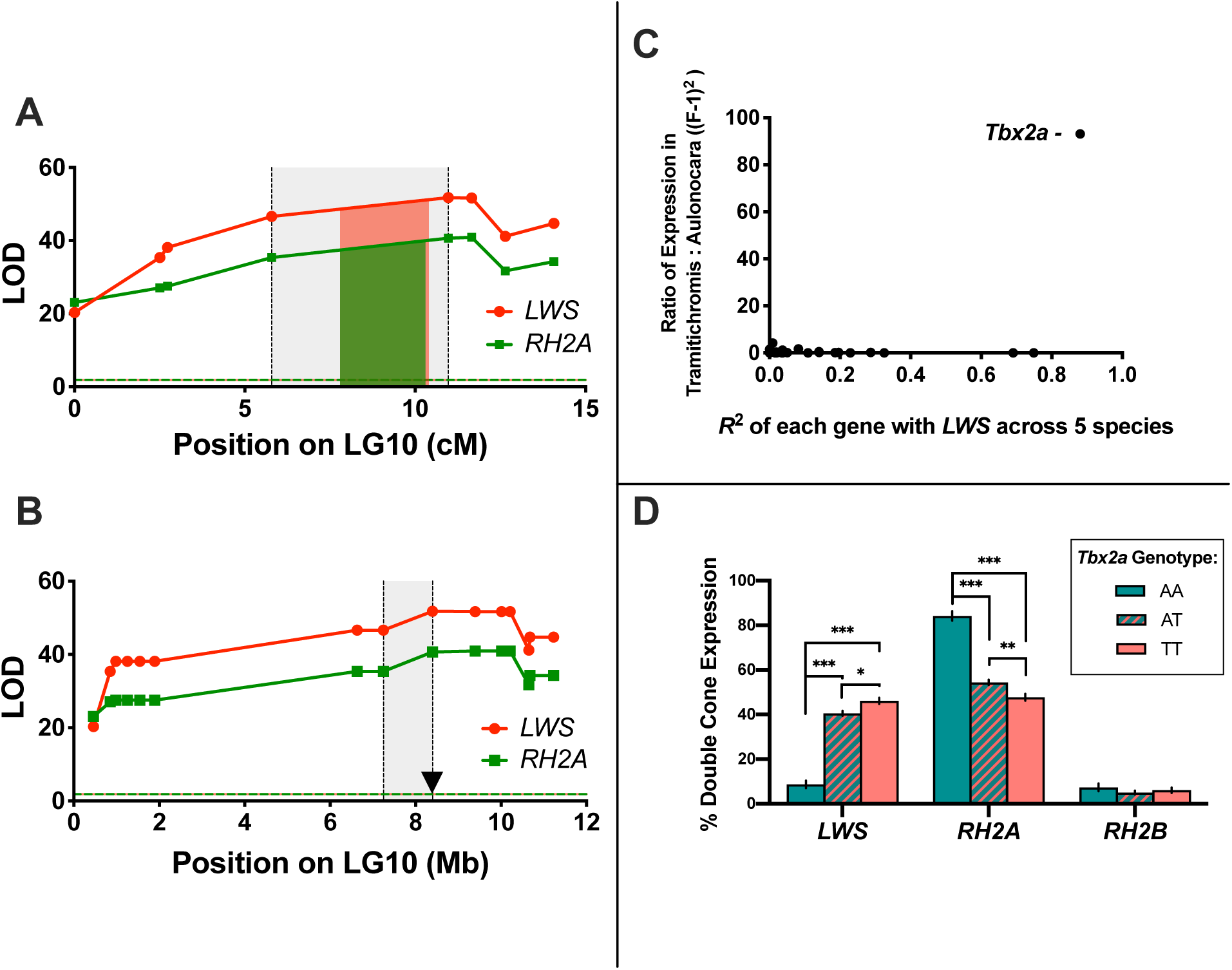
(A) Fine map of the *LWS* and *RH2A* eQTL from the cross between *A. baenschi* and *T. intermedius* in mapping space, and (B) markers mapped onto the *M. zebra* genome. Red and green regions denote the 95% Bayesian confidence interval for *LWS* and *RH2A* respectively. Grey shaded region indicates the space between the two markers surrounding the 95% confidence interval. The candidate gene *Tbx2a* is located within this region, roughly 8 kb from the highest associated microsatellite marker and denoted as a triangle. (C) Expression of the genes in the region between QTL markers surrounding the 95% confidence interval. Only the transcription factor Tbx2a is both highly correlated with *LWS* gene expression across species and strongly differentially expressed between *T. intermedius* and *A. baenschi*. X-axis: correlation between expression of each gene with *LWS* expression across five Malawi cichlid species not used in the cross. Y-axis: ratio of expression between *T. intermedius* (N=2) and *A. baenschi* (N=4) (given as: *(ratio-1)*^*2*^). (D) Opsin expression by *Tbx2a* genotype across 285 F_2_. AA= homozygous *A. baenschi Tbx2a*, TT= homozygous *T. intermedius Tbx2a*, and AT= heterozygous. *** P<0.0001, ** P<0.001, * P<0.01.

Given that the basic structure of the opsin regulatory network is conserved across closely related cichlid taxa, we examined previously sequenced retinal transcriptomes of five additional cichlid species as a second means of identifying transcription factors regulating *LWS*. Of these five taxa, three had high *LWS* expression and two had low/no *LWS* expression (19). Of the 31 eQTL genes, only three were strongly correlated (R^2^ > 0.7) with *LWS* expression (*ubc* (R^2^ = 0.92), *cenpv* (R^2^ = 0.74), and *Tbx2a* (R^2^ = 0.88)) (Supplementary Table 1).

*Tbx2a* was the only gene that was both differentially expressed between *T. intermedius* and *A. baenschi* and strongly correlated with *LWS* across the five additional cichlid species, suggesting that *Tbx2a* could explain the *LWS*/*RH2A* eQTL effect (Fig. 1C). Further support for this hypothesis was provided by the F_2_ from the cross as *LWS* and *RH2A* expression differed significantly between individuals that were homozygous for the *A. baenschi Tbx2a* and those with the *T. intermedius Tbx2a* genotype (Fig. 1D).

To confirm the correlation of *Tbx2a* with *LWS* expression, we first used qRT-PCR to measure retinal gene expression of *Tbx2a* and *LWS* relative to the cone specific gene *Gnat2*. We detected a strong correlation between *Tbx2a* and *LWS* expression across F_2_ offspring from our cross between *T. intermedius* and *A. baenschi* (R^2^=0.457, p<0.0001) (Fig. 2A). In addition, the expression of *Tbx2a* is strongly correlated with *LWS* expression across 58 additional species of Malawi cichlids that vary in *LWS* opsin gene expression (R^2^=0.847, p<0.0001) (Fig. 2B).

**Fig. 2.**
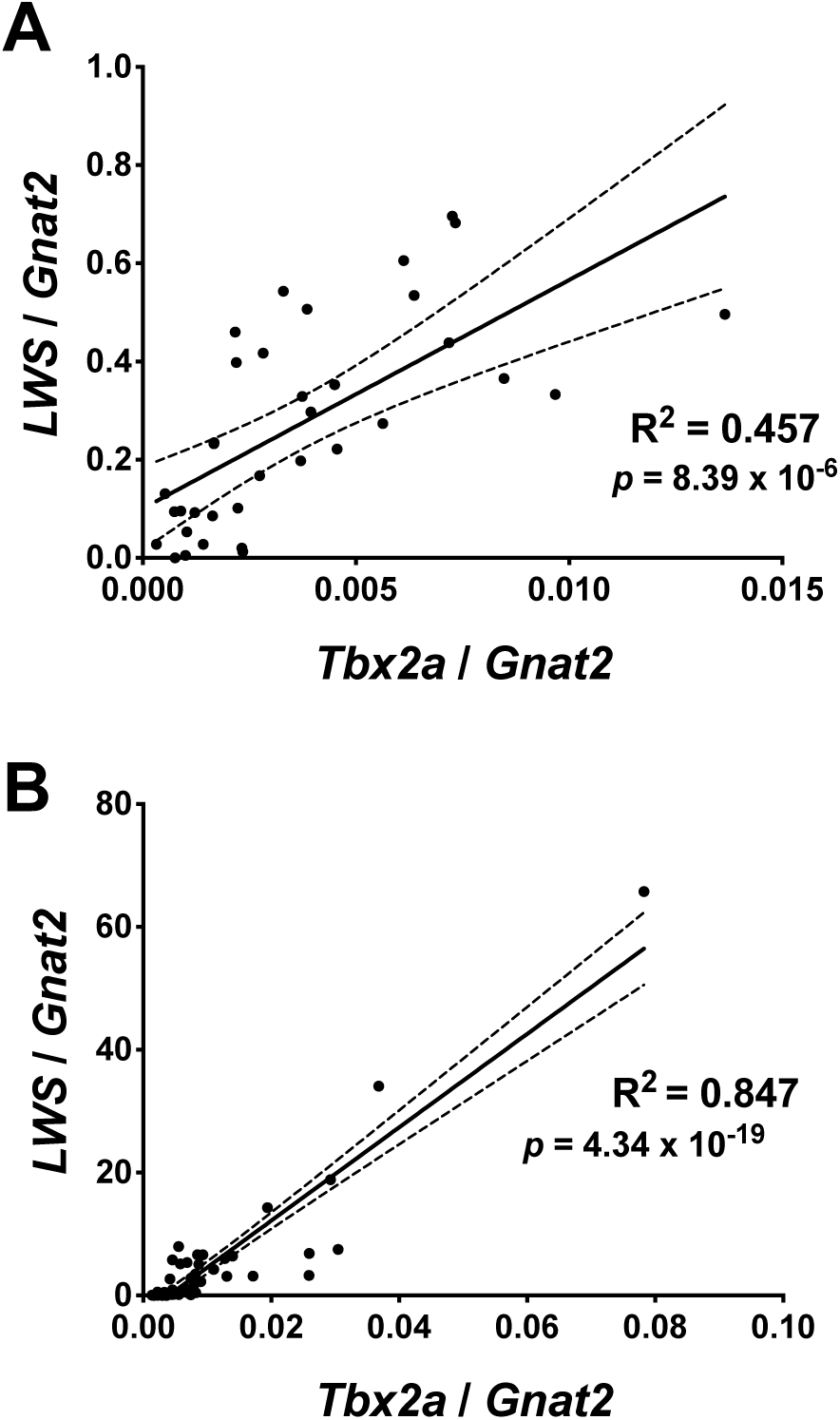
Retinal expression of *Tbx2a* and *LWS* is strongly correlated (A) across 35 *F*2 individuals from the cross between *T. intermedius* and *A. baenschi* in the lab, and (B) across 58 wild caught Malawi cichlid species. Solid line indicates the linear regression, dashed line denotes the 95% confidence interval of the regression and *p* value denotes the probability that the slope is non-zero.

The strong correlations of *Tbx2a* with *LWS* both within a cross and among natural species variants made *Tbx2a* a strong candidate to explain the *LWS*/*RH2A* eQTL. Furthermore, *Tbx2a* is located just 7.9 kb from the highest associated eQTL microsatellite marker, which explained 54.8% of the variance in *LWS* expression and 46.6% of variance in *RH2A* expression across 285 F_2_. We noted that members of the T-box 2 (Tbx2) transcription factor family can act as both activators and repressors of gene expression depending upon the promoter context and have been shown to play key roles in retinal development (20-22), with a reportedly prominent role for Tbx2b in specifying photoreceptor precursors to become short wavelength sensitive cone cells in zebrafish (23).

### How does Tbx2a regulate opsin expression?

In order to explore how Tbx2a could be modulating the switch in expression between *LWS*/*RH2A*, we identified transcription factor binding sites (TFBS) in regulatory regions of the *LWS* and *RH2A* opsins and tested the functionality of these sites experimentally. In cichlids, there are two *RH2A* loci (*RH2Aα* and *RH2Aβ*) that are functionally identical but have unique promoter sequences. To identify possible Tbx2a binding sites, we compared 800 bp of the proximal promoter from each of *RH2Aα* and *RH2Aβ* to the *H. sapiens* TBX2 transcription-factor binding matrix (JASPAR database of transcription factor binding profiles). We could identify only one TBX2 binding site upstream of the *RH2Aβ* transcriptional start site (TSS) (relative score = 0.921) and none upstream of the *RH2Aα* TSS.

The *RH2* gene array is downstream of a Locus Control Region (LCR) that controls *RH2* gene expression in zebrafish (24, 25). To identify whether Tbx2a could be interacting with the RH2 LCR, we aligned the RH2 LCR of *A. baenschi, T. intermedius*, carp (*Cyprinus carpio*), zebrafish (*Danio rerio*), stickleback (*Gasterosteus aculeatus*), and medaka (*Oryzias latipes*) (Fig. 3). We detected a 16 bp region that was perfectly conserved across all six species (representing ∼230 Mya divergence) and contained a strongly-predicted Tbx2 binding site based on the *H. sapiens* TBX2 matrix (relative profile score of 0.97). To test whether the cichlid Tbx2a transcription factor binds to the predicted sites in the RH2 LCR or *RH2Aβ* promoter, we cloned the coding region of *Tbx2a* into the pcDNA4-HisMax C vector and expressed the recombinant cichlid Tbx2a in the mammalian cell line MDCK (Madin-Darby Canine Kidney). We used the Tbx2a enriched nuclear fraction for Electrophoretic Mobility Shift Assays (EMSA) with probes containing the predicted Tbx2a binding sites in the RH2 LCR and *RH2Aβ* promoter. The cichlid Tbx2a did not bind to the *RH2Aβ* promoter probe but shifted the RH2 LCR probe with high affinity. Furthermore, cichlid Tbx2a did not bind to a mutant version of the probe where the four bases corresponding to the key bases in the *H. sapiens* Tbx2 binding motif were changed (Fig. 4A; Supplementary Table 2). These results show that the cichlid Tbx2a can bind to the highly conserved TBX2 TFBS in the RH2 LCR.

**Fig. 3.**
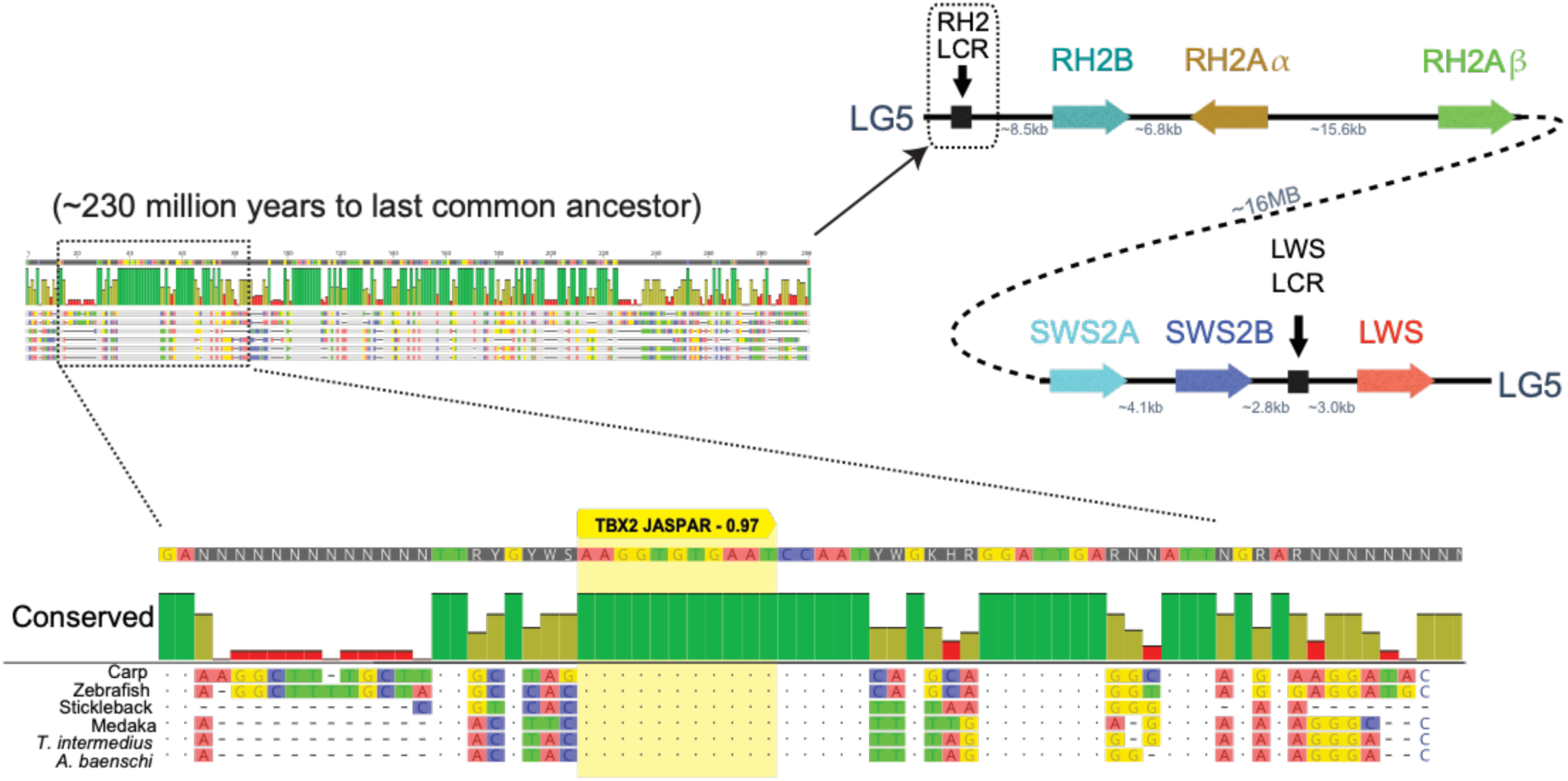
The RH2 Locus Control Region (LCR) identified in zebrafish is highly conserved across teleosts. A Tbx2 binding site (yellow) is strongly predicted in the longest conserved region of the RH2 LCR when compared to the JASPAR database (score 0.97).

**Fig. 4.**
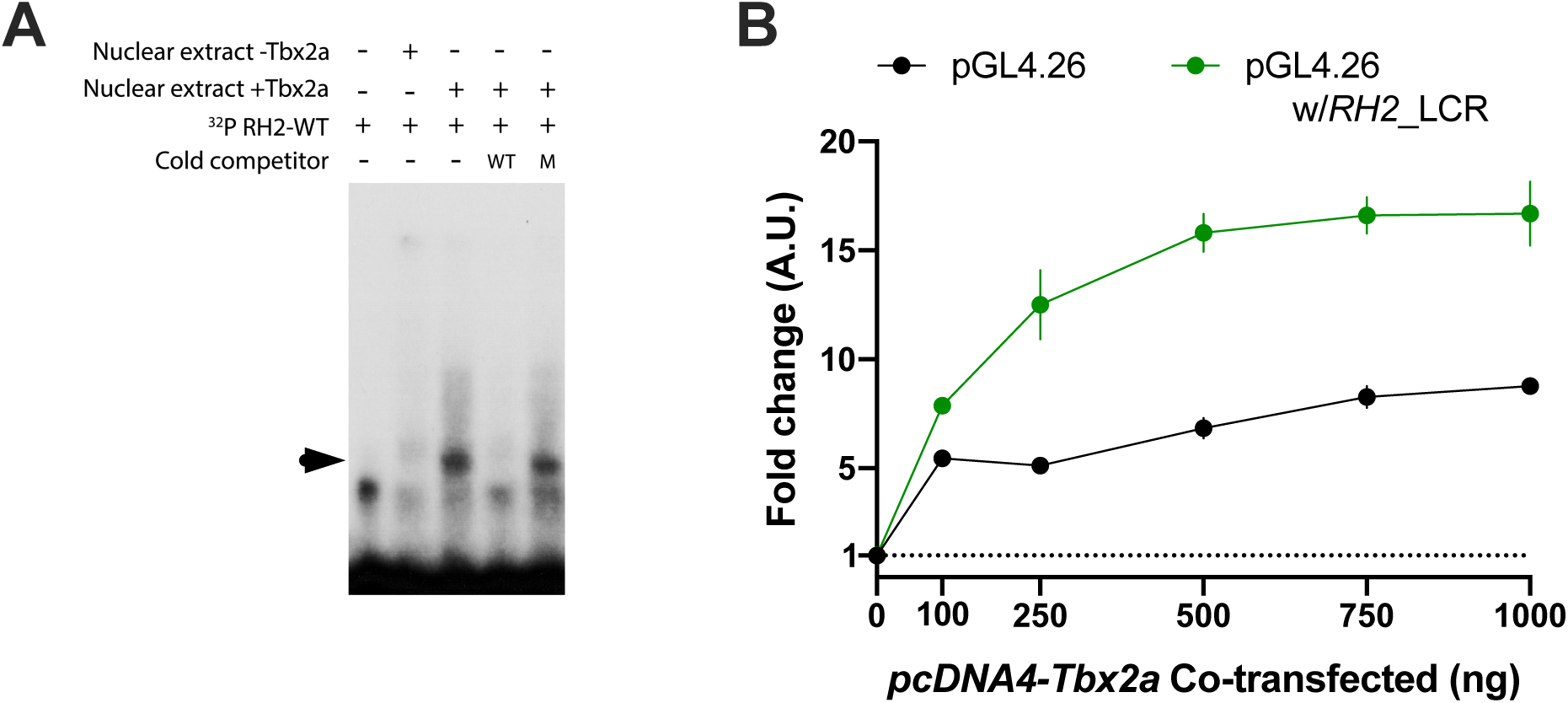
(A) Tbx2a directly binds to RH2 LCR in vitro. Radioactive-labeled RH2 LCR probes containing the putative Tbx2a binding site were incubated with nuclear extracts of MDCK cells transfected with empty vector (-Tbx2a) and vector encoding the cichlid Tbx2a transcription factor (+Tbx2a). The shift created by protein-DNA binding (indicated by arrowhead) was reduced by the addition of unlabeled probe in 400x molar excess (cold wild-type competitor, WT), but to a lesser degree by unlabeled probes with critical mutations in binding site residues (cold mutant competitor, M). (B) MDCK cells were co-transfected with constructs containing the RH2 LCR sequence upstream of a minimal promoter driving firefly luciferase reporter gene with increasing concentrations of Tbx2a expression plasmids. Constructs without the RH2 LCR sequence were used as a control. Fold change is relative to reporter gene activation in the absence of Tbx2a expression vectors.

To test whether Tbx2a binding to the RH2 LCR affects downstream gene expression, we cloned the RH2 LCR into a pGL4.26 vector upstream of luciferase driven by a minimal promoter. Luciferase expression increased substantially with increasing amounts of Tbx2a during co-transfection in MDCK cells but only in the presence of the pGL4.26 plasmid containing the RH2 LCR (Fig. 4B). We detected a similar pattern of expression in both IMCD3 and HEK293 cell lines (Supplementary Fig. 1). These results demonstrate that the interaction between Tbx2a and the RH2 LCR is capable of modulating downstream gene expression.

The *LWS* opsin gene is also regulated by an LCR and a proximal promoter. The LCR is located ∼3 kb upstream of the *LWS* coding sequence (26). JASPAR predicted two TBX2 TFBS in the *LWS* LCR but with lower confidence (relative scores of 0.85 and 0.81) than the site in the *RH2* LCR (score of 0.97). Our EMSA results suggest that cichlid Tbx2a does not bind to either of these sites. JASPAR also predicted two TBX2 TFBS with a relative score of ≥ 0.90 (0.92 located 1,178 bp upstream and 0.96 located 471 bp upstream of the *LWS* TSS) within the 3 kb region between the *LWS* LCR and the beginning of the *LWS* coding sequence. EMSA demonstrated the binding of cichlid Tbx2a to the site 471 bp upstream of *LWS* TSS but not to the 1,178 bp upstream site (Fig. 5A).

**Fig. 5.**
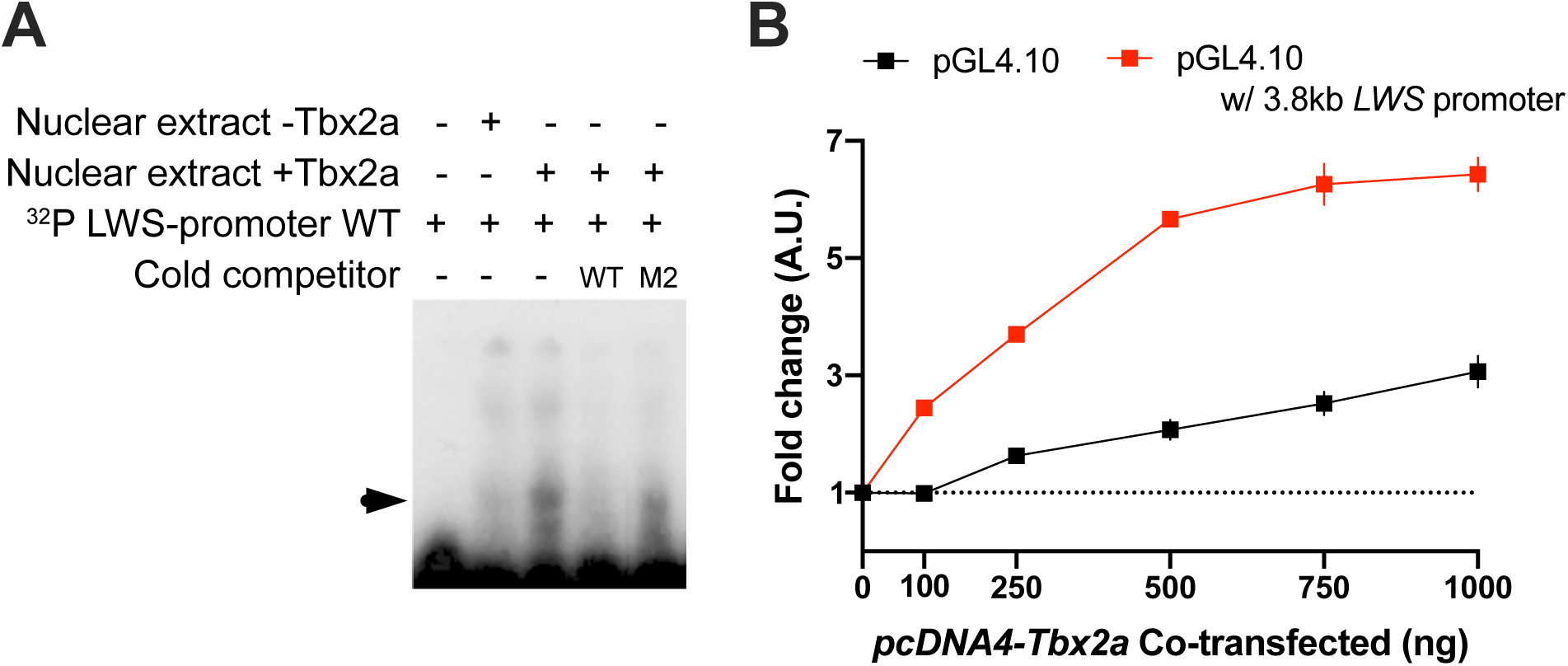
(A) Tbx2a directly binds to *LWS* promoter in vitro. Radioactive-labeled *LWS* promoter probes containing the putative Tbx2a binding site were incubated with nuclear extracts of MDCK cells transfected with empty vector (-Tbx2a) and vector encoding the cichlid Tbx2a transcription factor (+Tbx2a). The shift created by protein-DNA binding (indicated by arrowhead) was reduced by the addition of unlabeled probe in 400x molar excess (cold wild-type competitor, WT), but to a lesser degree by unlabeled probes with critical mutations in binding site residues (cold mutant competitor, M). (B) MDCK cells were co-transfected with constructs containing the *LWS* promoter sequence upstream of a firefly luciferase reporter gene with increasing concentrations of Tbx2a expression plasmids. Constructs without the *LWS* promoter sequence were used as a control. Fold change is relative to reporter gene activation in the absence of Tbx2a expression vectors.

To test whether Tbx2a binding to the *LWS* promoter can modulate gene expression, we cloned the full 3.8 kb from upstream of the *LWS* LCR to the *LWS* TSS into a pGL4.10 vector upstream of a promoter-less copy of the luciferase gene. Tbx2a showed a much larger increase in luciferase expression when transfected with pGL4.10 vector containing the upstream *LWS* region compared to the control plasmids (Fig. 5B). Thus, the interaction between Tbx2a and the *LWS* regulatory region is capable of modulating gene expression.

### How has Tbx2a driven differences across species?

To identify how Tbx2a might lead to the *LWS*/*RH2A* shift in visual tuning across species, we asked if differences in *Tbx2a* expression across species is due to changes in linked regulatory sequence between *T. intermedius* and *A. baenschi*. We used transcriptomes from *T. intermedius, A. baenschi* and four F_1_ from the cross to examine allele specific expression and determined that 85.6% of the variance in *Tbx2a* expression is due to differences in *cis*-linked regulatory sequence. Our results strongly suggest that a change in regulatory sequence linked to the *Tbx2a* locus can explain the *LWS*/*RH2A* eQTL.

To identify the regulatory variant, we sequenced whole genomes from *T. intermedius* (N=2) and *A. baenschi* (N=2) and mapped the reads onto the *M. zebra* reference genome (coverage of 13.13 ± 9.19, 13.02 ± 8.99, 11.04 ± 7.71, and 9.13 ± 6.58, respectively). Our analyses uncovered only 10 fixed variants between *T. intermedius* and *A. baenschi* from the start of *Tbx2a* to the next gene upstream (protein phosphatase 1D) within a region of ∼19.5 kb, and only three fixed differences were identified from the end of *Tbx2a* to the next gene downstream (acetyl-CoA carboxylase alpha) (∼5.5 kb) (Fig. 6, Supplementary Table 3). The largest of these differences was a 938 bp deletion in *A. baenschi* located 13.4 kb upstream of the *Tbx2a* start site. One other deletion downstream of *Tbx2a* in *A. baenschi* was only one bp in length. The remaining 11 fixed differences were single base pair changes. Interestingly, the 968 bp deletion contained a Mariner-8 transposon, a ‘cut-and-paste’ transposable element (27).

**Fig. 6.**
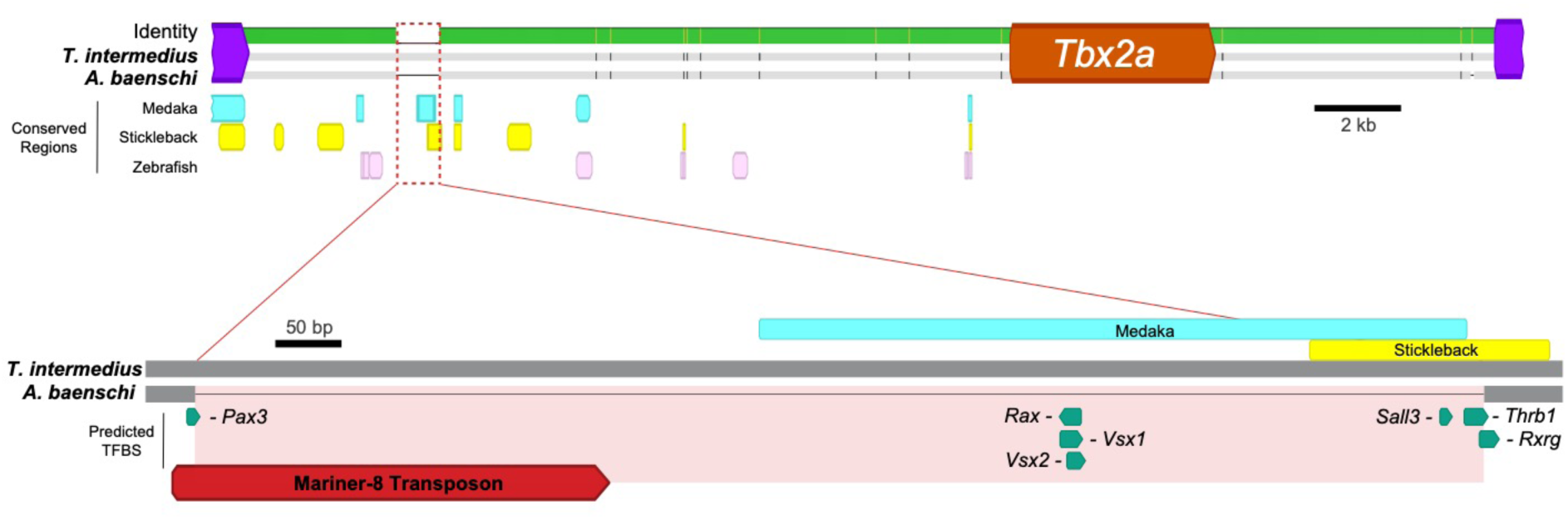
Top: Alignment of the *Tbx2a* region between *T. intermedius* and *A. baenschi* from the next gene upstream (protein phosphatase 1D) to the next gene downstream (acetyl-CoA carboxylase alpha). The 13 differences between species are denoted as vertical black lines (for single base pair differences) or horizontal lines (for sequence missing in one species). Region identified by *MultiPipMaker* to be conserved are shown as colored rectangles for medaka (blue), stickleback (yellow) and zebrafish (pink). Bottom: The 968 bp that is deleted in *Aulonocara baenschi* (shaded red). This region shows some conservation with both Stickleback and Medaka and contains several predicted TFBS for important retinal transcription factors as well as Pax3 (known to regulate Tbx2). The presence of a Mariner-8 transposon suggests this region was lost when this transposon excised from the area and took important regulatory sequence with it.

To delineate regulatory implications of sequence differences, we undertook a phylogenomic approach and identified intergenic sequences that are conserved between *T. intermedius*, stickleback *(Gasterosteus aculeatus*) (∼128 Mya divergence), medaka (*Oryzias latipes*) (∼119 Mya divergence), and/or zebrafish (*Danio rerio*) (∼230 Mya divergence). Of the 18,818 bp between *Tbx2a* and the next gene upstream in *T. intermedius*, the *MultiPipMaker* software identified only 4.6% of the sequence that was conserved between *T. intermedius*, stickleback and medaka, 1.8% conserved between *T. intermedius*, medaka and zebrafish, and only 0.3% was conserved among all four. The 938 bp deleted in *A. baenschi* contained the sequence that was partially conserved in both stickleback and medaka (Fig. 6). In this region we also identified predicted binding sites for six transcription factors that are known to play key roles in retinal development across vertebrates (28, 29); *Rxrg, Thrb, Sall3, Vsx1, Rax*, and *Vsx2* (Fig. 6). Moreover, the cichlid sequence contained a predicted binding site for *Pax3* which is known to regulate *Tbx2* expression (30). All of these potentially important regulatory sites have been lost in *A. baenschi*, which may explain the difference in *Tbx2a* expression between *T. intermedius* and *A. baenschi*.

## DISCUSSION

Spectral tuning of color vision is frequently accomplished by rapid evolutionary changes in opsin gene expression (9, 13, 31). However, evolution of gene expression through multiple mutations of independent gene regulatory elements would be inherently slow since two simultaneous changes in opposite directions are not very likely (32). We propose that this could be resolved by regulatory elements that simultaneously downregulate one opsin while upregulating another opsin. Thus, a switch requiring only a single change could permit the rapid evolutionary changes in visual tuning across close taxa. Here, we show that the transcription factor Tbx2a modulates an established switch between *RH2A* and *LWS* opsin gene expression. Using DNA-binding proteins and luciferase reporter assays, we show that Tbx2a can directly interact with the RH2 LCR and *LWS* promoter regions. We propose that a single change in the regulatory region upstream of *Tbx2a* is likely responsible for the rapid shift in visual tuning between *T. intermedius* and *A. baenschi*. This work therefore provides strong support for opsin regulatory elements that control multiple opsin genes.

### Tbx2a is a major regulatory component of switching between RH2A and LWS opsins

We have identified Tbx2a as a key component in modulating the switch between *RH2A* and *LWS* opsin gene expression by fine mapping an eQTL on LG10 in a cross between *Aulonocara baenschi* (no *LWS* expression) and *Tramitichromis intermedius* (high *LWS* expression). Support for the idea that *Tbx2a* explains the eQTL comes from its location only ∼8 kb from the highest associated microsatellite marker and the strong correlation of *Tbx2a* expression with *LWS* opsin expression in *F*2 hybrids. *Tbx2a* expression was also strongly correlated with *LWS* expression across 58 other species of Lake Malawi cichlids, as well as four cichlids in other African lakes and rivers, indicating that the role of *Tbx2a* is largely conserved across cichlids.

Many of the genes involved in photoreceptor gene regulation are highly conserved from fish to mammals (33). The Tbx2 class of transcription factors predates the vertebrate lineage and unlike the other T-box transcription factors, members of the Tbx2/3 class are known to act as repressors as well as activators (34). Musser and Arendt (28) showed that the Tbx2 class has played a strong role in photoreceptor development since well before the emergence of the mammalian lineage. While mammals have lost the RH2 class of opsins, they have maintained both LWS and TBX2. This raises the possibility that TBX2 may be acting as a conserved transcription factor modulating *LWS* expression. Nearly 20 years ago *Tbx2* was found to be expressed along the same dorsal-ventral pattern as *M*/*Lws* opsin during development in the mouse retina, which has high *M/Lws* expression in the dorsal, but not ventral, retina (35). More recently, this possibility was further supported by a dataset from single cell transcriptomes of mouse retinal cells generated by Macosko *et al.* (36). Reanalysis of this dataset revealed that none of the “pure” *Sws1* cones expressed *Tbx2*, yet some of the cones co-expressing *Sws1* and *Lws* did show *Tbx2* expression (28, 36). This pattern raises the possibility that the role of *Tbx2a* in promoting *LWS* expression could be conserved in even distantly related vertebrate lineages.

### How does Tbx2a regulate opsin expression?

To determine how Tbx2a is modulating *RH2A* and *LWS* opsin expression, we examined predicted Tbx2 binding sites located in *RH2A* and *LWS* opsin regulatory sequence. Within the RH2 LCR we found a conserved Tbx2a predicted-binding site that was completely conserved across six taxa (representing ∼230Mya). Such strong conservation suggests this location plays an important role in the mechanism by which the LCR controls downstream *RH2* gene expression. Our EMSA results revealed that the cichlid Tbx2a does indeed bind to the RH2 LCR at this location and our luciferase results show that this Tbx2a-RH2 LCR complex can dramatically change downstream gene expression. Since the interaction between Tbx2a and the RH2 LCR occurs at a highly conserved TFBS, it suggests an important role for Tbx2a in mediating how the RH2 LCR operates, which should be further explored.

Similar results were obtained with the *LWS* promoter. Our EMSA tests revealed that Tbx2a can indeed bind to a TFBS located 471 bp upstream of the *LWS* transcription start site and this Tbx2a-*LWS* promoter complex can dramatically change downstream gene expression.

It was unexpected for Tbx2a to increase expression of luciferase downstream of both the *LWS* promoter and the RH2 LCR. In cichlid retina, *Tbx2a* is directly correlated with *LWS* expression, but inversely correlated with expression of *RH2A*. However, Locus Control Regions are generally controlled by multiple transcription factors that interact with one another (37). It seems plausible that Tbx2a requires additional cichlid cone-cell specific co-factors to produce the anticipated effect of decreasing *RH2A* expression that are not present in the cell lines used in our luciferase experiments. Our recent QTL effort involving a new cichlid cross has identified additional genetic factors that contribute to variation in *RH2A* expression (38) and these may help explain the cell specific actions of the RH2 LCR on *RH2A* expression.

### Single changes to opsin regulatory elements can switch visual tuning between species

Allele specific expression in F_1_ hybrids between *A. baenschi* and *T. intermedius* confirmed a *cis*-regulatory change in *A. baenschi* underlying the decrease in *Tbx2a* expression. Comparing differences in the predicted TFBS for the fixed differences between *A. baenschi* and *T. intermedius* around the *Tbx2a* gene revealed that the only region to contain predicted TFBS for retinal transcription factors was a 968 bp deletion located 13 kb upstream of *Tbx2a* in *A. baenschi*. Within this 968 bp deletion, 119 bp was conserved between *T. intermedius*, stickleback and medaka. Conserved non-coding sequences often indicate important regulatory function. In this region, we found predicted TFBS for known retinal transcription factors including members of the *Thrb* and *Rx* families. In medaka, knocking out *Rx3* resulted in the absence of *Tbx2* expression in the retina, and plasmid injections of *Rx3* rescued *Tbx2* expression (39), suggesting a feedback link between *Rx* and *Tbx* genes. Thus, this 968 bp deletion in *A. baenschi* might have resulted in the difference in *Tbx2a* expression between *A. baenschi* and *T. intermedius*.

Within the 968 bp deletion, we also detected a Mariner-8 transposable element (TE). The mariner class of TEs make a double strand break as they move throughout the genome and when the break is repaired through single-strand annealing, a deletion often occurs in the DNA flanking the break (40). African cichlid fishes are known to have high rates of TE movement (19, 41-43). We hypothesize that the deletion in *A. baenschi* was caused by a Mariner-8 transposable element leaving the area and taking the important *Tbx2a* regulatory sequence with it.

## CONCLUSION

We have identified an eQTL variant that allows for simultaneous upregulation of one opsin and downregulation of another. We identified the transcription factor Tbx2a as a major component of the regulatory framework, which drives this switch between *RH2A* and *LWS* opsin genes by binding to both the RH2 LCR and the *LWS* promoter. Our data demonstrate that single mutations within the components of this network can result in instant changes in visual tuning by changing the expression of multiple genes, rather than independent changes in the regulatory regions of each opsin gene. One possible explanation for this switch is that it evolved to facilitate developmental shifts in opsin expression. This would enable the single mutation observed here to cause heterochronic shifts in the developmental pathway and contribute to rapid evolutionary change. Future work should consider how such developmental switches evolve.

## METHODS

### Fine mapping the LWS/RH2A opsin eQTL on LG10

In our previous study of opsin eQTL we established a cross between *T. intermedius* and *A. baenschi* and used restriction-site associated DNA sequences (RAD-seq) across 115 F_2_ to identify overlapping eQTL on LG10 that explained 48.6% and 57.05% of the expression of *LWS* and *RH2A* respectively (17). Here we developed eight fluorescently-labeled microsatellite markers across these eQTL and genotyped 288 F_2_ adults. Primers flanking the microsatellites were designed with Primer3 (44) and obtained from Eurofins MWG Operon (Huntsville, AL). Forward primers were either fluorescently labeled, or ordered with a CAG tag (GCAGTCGGGCGTCATCA) that matched a fluorescent CAG primer which was labeled with either 6’FAM or HEX dye. PCR products were run on an ABI 3730xl Genetic Analyzer and marker genotype was determined in GeneMapper® (ABI). All individuals were sacrificed with an overdose of MS-222; DNA was extracted from fin clips using a DNeasy Kit (Qiagen) and retinas were dissected and stored in RNAlater (Invitrogen) at −20°C prior to RNA extraction.

Opsin expression phenotypes were determined by quantitative real-time PCR (qRT-PCR). Total RNA was extracted from whole retinas using a RNeasy kit (Qiagen) including a DNase step, and 0.5 µg was used to make cDNA with SuperScript III (Invitrogen) following manufacturer’s protocol. qRT-PCR was conducted for each of the cone opsins on a Roche 480 Lightcycler as described previously (45). Briefly, the seven cone opsins present in East African cichlids were measured individually using custom Taqman® assays (Invitrogen) with the exception of *RH2Aα* and *RH2Aβ*, which were measured in combination because the sequence similarity between these genes is too high to tell them apart with qPCR-based methods (46). Results were corrected for relative assay efficiencies calculated by running a multi-opsin construct containing all targets in a 1:1 ratio and the absolute efficiency of *RH2A* was determined from serial dilutions of cDNA (47).

The ratio of single to double cones in the cichlid retina is fixed in a highly structured retinal mosaic (48). *SWS* genes are only expressed in single cones, while *RH2* and *LWS* genes are only expressed in double cones (48, 49). Therefore, we represent the expression of each of the double cone opsins as:

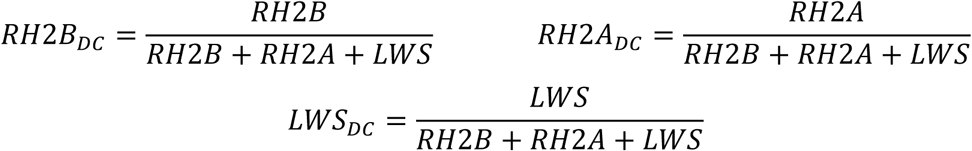

The eQTL was fine mapped by combining the previous RAD-seq markers with fluorescent microsatellite markers to provide 15 markers spanning the previously identified *LWS*/*RH2A* eQTL. Marker order was identified by BLASTN (version 2.2.28+) (50, 51) against the *Metriaclima zebra* reference genome (41). eQTLs and 95% Bayesian confidence intervals (CI) were identified for *LWS* and *RH2A* using R/qtl (52) and the percent of variance in *LWS* and *RH2A* expression that was explained by the closest marker to *Tbx2a* was determined using the *lm* function in R v3.5.2 (53). We conservatively identified putative transcription factors by finding all genes in the *M. zebra* UMD2a reference genome (41) between the two markers flanking the 95% CI (corresponding to a 1.15 Mb region).

### Retinal transcriptome sequencing, analyses, and additional species

To facilitate transcriptome comparisons across species we first mapped the NCBI Refseq genes for *Oreochromis niloticus* (RefSeq #GCF_001858045) onto the *M. zebra* genome (41) using GMAP (version 2015-07-23) (54) to annotate the corresponding set of genes in *M. zebra*. We then generated retinal transcriptomes from four *A. baenschi* and two *T. intermedius* using a TruSeq RNA sample preparation kit v.2 (Illumina) and run on an Illumina HiSeq1500 with 100 bp single-end reads. The transcriptome reads were aligned to the gene models predicted in the *M. zebra* genome. *Cuffdiff* (version 2.2.1) was used to quantify transcript expression in terms of FPKM (Fragments per Kilobase of gene model per Million reads), which corrects for the size of the gene and the number of reads per sample (55). To quantify differential expression between *T. intermedius* (T) and *A. baenschi* (A) we followed Schulte *et al*. (2014) and took whichever ratio of FPKM was greater than 1 (*F* = T/A or A/T), subtracted 1 and squared [(*F*-1)^2^]. This provides a measure that is close to zero for genes with similar expression and a larger value for genes which are more highly differentially expressed in one species relative to the other.

We then compared differential expression between *T. intermedius* and *A. baenschi* to retinal transcriptomes of five other East African cichlids that were generated as part of the Broad Institute’s Cichlid Genome Project (19). Two of the species express no *LWS*: *M. zebra* (Broad assembly v.0), and *Neolamprologus brichardi* (Broad assembly v.1). Three express high *LWS*: *Pundamilia nyererei* (Broad assembly v.1), *Astatotilapia burtoni* (Broad assembly v.1), and *Oreochromis niloticus* (Broad anchored assembly v1.1). To compare genes between species we mapped the NCBI RefSeq transcripts for *Oreochromis niloticus* (RefSeq #GCF_001858045) onto the genome for each of the five species using GMAP (54). We then aligned retinal transcriptome reads and quantified relative gene expression in each species as FPKM using Tophat2 and *Cuffdiff* (55).

Of the 31 genes in the 95% C.I. two were only expressed in one species and excluded from correlation analyses; the correlation of expression with *LWS* opsin expression (FPKM) was calculated using linear regressions across the five species. To test whether transcriptomes provided reliable indicators of gene expression, we compared the relative opsin expression levels within the transcriptomes to our previous qRT-PCR studies and found there to be good agreement (56).

### Identifying ‘Tbx2-like’ as Tbx2a

We found the gene annotated in the *M. zebra* genome as ‘*Tbx2-like*’ to be both highly correlated with *LWS* across species and highly differentially expressed between *T. intermedius* and *A. baenschi*. To determine the true identity of this ‘*Tbx2-like*’ locus we built a phylogeny including *Tbx2* sequences of eight additional vertebrate species (*Danio rerio, Gasterosteus aculeatus, Oryzias latipes, Oreochromis niloticus, Gallus gallus, Homo sapiens, Bos taurus*, and *Mus musculus*). Sequences were aligned using MAFFT (57) and maximum likelihood trees were constructed using GARLI (58) with 100 bootstrap replicates to determine branch support (Supplementary Fig. 2). The ‘*Tbx2-like*’ fell within the *Tbx2a* clade, and identity was further confirmed by synteny in the UCSC genome browser across five species (*M. zebra, O. niloticus, O. latipes, G. aculeatus* and *D. rerio*) (Supplementary Fig. 3).

### Genome sequencing of T. intermedius and A. baenschi

Genomes from two individuals of *T. intermedius* and two individuals of *A. baenschi* were sequenced using a TruSeq DNA sample preparation kit and run on an Illumina HiSeq1500 (100 bp paired-end reads with 550 bp inserts). Reads were filtered then quality checked and trimmed with Trimmomatic (59) before being aligned to the *M. zebra* reference genome (41) using *bwa-mem* (version 0.7.12-r1039) (60). Alignments were converted to BAM format, duplicates were marked and whole genome metrics were determined using Picard (version2.1.0). Alignments were viewed in IGV (61) and regions of interest were edited manually in Geneious 11 before comparing the consensus sequence between species.

### Correlating expression of Tbx2a and LWS

The expression of *Tbx2a, LWS*, and *Gnat2* (a cone-specific gene) was measured from retinal cDNA using custom probe based quantitative real-time PCR (qPCR) assays (Taqman® probes (Invitrogen)) (See Supplementary Table 4 for primer/probe sequences and efficiencies). Assay efficiencies were calculated from a 1000-fold dilution series of a custom gene construct that contained the five target opsin genes in a 1:1 ratio. The expression of *LWS* and *Tbx2a* in each individual was normalized to expression of *Gnat2* using the equation:

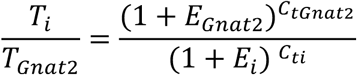

such that *E* is the assay efficiency found from the dilution series and *Ct* is the critical threshold of that reaction in the sample. To determine if expression of *Tbx2a* is correlated with *LWS* expression across species we conducted qPCR on 58 wild caught Malawi cichlid species (see Supplementary Table 5 for species list and (45) for description of collections). To determine the strength of the correlation of *Tbx2a* with *LWS* within the cross between *A. baenschi* and *T. intermedius* we conducted qPCR on 35 F_2_ individuals. We ran a linear regression between the two genes in each dataset in GraphPad Prism v.8.0.1.

### Cis versus trans control of Tbx2a expression using allele specific expression

To determine if differences in *Tbx2a* expression were due to linked differences in regulatory sequence (thus supporting *Tbx2a* as a candidate for driving the *LWS*/*RH2A* eQTL) we identified SNPs in the expressed *Tbx2a* sequence that were differentially fixed between *A. baenschi* and *T. intermedius* using the transcriptomes described above. We then generated retinal transcriptomes from four F_1_ from the cross using a TruSeq RNA Sample preparation kit v.2 (Illumina) and sequenced on an Illumina HiSeq1000 with 100 bp single-end reads. Reads were mapped onto the *M. zebra* genome as before. We compared the number of reads of *T. intermedius* alleles to the number of reads of *A. baenschi* alleles. We then compared the log2 of this ratio to the log of the ratio of expression between the *A. baenschi* and *T. intermedius* parents (measured as FPKM-described above).

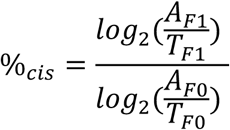

Values closer to 1 reveal more of the variance in *Tbx2a* expression is explained by *cis*-linked regulatory differences (62).

### Identifying fixed variants between species linked to Tbx2a

The genomes of *T. intermedius* (N=2) and *A. baenschi* (N=2) (described above) were compared to identify fixed differences around *Tbx2a* from the next gene upstream (Protein Phosphatase 1D (XM_004557358), located 19.5 kb upstream) and the next gene downstream (Acetyl-CoA carboxylase alpha (XM_014408615), located 5.5 kb downstream) in Geneious v.11. We found one large deletion in *A. baenschi* and verified this deletion with PCR and Sanger sequencing that spanned the deletion. To verify that this deletion was not specific to our lab population we obtained Sanger sequence reads that spanned the deletion from two unrelated *A. baenschi* individuals collected in the wild in 2012 from Nkhoma Reef, Lake Malawi, Africa.

### Transposable element within A. baenschi deletion

To search for transposable elements in the vicinity of the *A. baenschi* deletion we compared the intact sequence from *T. intermedius*, including 100 bp before and after the deletion, to the GIRI Repbase repository of transposable elements using CENSOR (63).

### Genomic footprinting to identify sequence conservation upstream of Tbx2a

As *Tbx2a* is an evolutionarily conserved gene; important regulatory sequence is likely to be conserved across distantly related species. To find such conserved regions upstream of *Tbx2a* we gathered the sequence between *Tbx2a* and the next gene upstream (protein phosphatase 1D (PPM1D)) from the medaka (23 kb), zebrafish (401 kb), and stickleback (18 kb) reference genomes on the UCSC genome browser. We then identified regions of conservation between these species and *T. intermedius* using MultiPipMaker (64).

### Identifying transcription factor binding sites (TFBS)

The TRANSFAC repository of vertebrate transcription factor binding sites (65) was used to predict TFBS at *p* < 0.0001 from the *T. intermedius* sequence that was missing in *A. baenschi*. The list was then reviewed for transcription factors known to play key roles in vertebrate photoreceptor development (28).

### Tbx2 binding sites in RH2 and LWS locus control regions

The RH2 and LWS classes of opsins each have their own locus control regions (LCR) that have been highly conserved across a wide range of taxa (25, 26, 66). These LCRs match between *A. baenschi* and *T. intermedius*. Using the JASPAR database, we scanned for Tbx2 binding sites (relative score >80%) in the RH2 and LWS LCRs. We also scanned for Tbx2 binding sites (relative score >90%) in the 3 kb region between *LWS* coding region and the LWS LCR, and 800 bp upstream of the *RH2Aα* and *RH2Aβ* genes.

### Generating cichlid Tbx2a protein

A gBlock^®^ synthetic gene from Integrated DNA Technologies was designed to contain the full 1,839 bp of cichlid *Tbx2a* coding sequence flanked by restriction sites. To clone this gBlock® into the pcDNA™4-HisMax C mammalian expression vector (ThermoFisher Scientific) both were double digested (*KpnI*-HF and *XhoI*) and run on a PCR clean up column (Machery-Nagel). The cut products were ligated together with a Quick Ligation™ kit (NEB) and used to transform DH5α *E. coli*. Colonies were PCR screened for presence of the correct insert size. Correct reading frame and coding sequence of the construct was verified by Sanger sequencing. The pcDNA™4-HisMax vector adds both poly-histidine and Xpress™ tags to the N terminal of expressed proteins. Because there is no cichlid-specific Tbx2a antibody, α-Xpress antibody was used for all western blots.

To produce recombinant Tbx2a protein, Madin-Darby Canine Kidney (MDCK) cells were transfected with the pcDNA4-*Tbx2a* construct using Lipofectamine 3000 (ThermoFisher Scientific) following manufacturer’s protocol. Cells were harvested 48 hours after transfection and nuclear proteins were obtained with the classical method of nuclei isolation in hypotonic buffer. Protein extracts were quantified by BCA protein assay according to manufacturer’s protocol (Pierce) and solubilized in 4x Laemmli buffer + β-mercaptoethanol. After denaturation, 7 μg of nuclear protein extract was separated by SDS-PAGE and transferred to PVDF membrane (Trans-Blot Turbo System, Bio-Rad). We followed standard immunoblot procedures using primary antibodies raised against the Xpress tag (ThermoFisher R910-25, 1:5,000), and anti-mouse IgG HRP-conjugated secondary antibodies (Jackson ImmunoResearch 715-035-150, 1:10,000). Detection was performed by enhanced chemiluminescence (ECL) using the SuperSignal West Pico Plus system (ThermoFisher). The expressed protein matched the predicted size for the cichlid Tbx2a (72 kDa) and was localized to the nucleus (Supplementary Fig. 4).

### Electrophoretic Mobility Shift Assay (EMSA)

The predicted binding motif of Tbx2 in *H. sapiens* is 11 bases long. We designed all EMSA probes to also include 15 bases upstream and downstream of the predicted Tbx2 binding site, making all probes 41bp in length (Supplementary Table 2). Mutant probes were designed such that the most influential sites in the Tbx2 binding motif were changed to intolerable bases. Probes were generated by annealing complementary oligos in duplex buffer (10 mM Tris-HCl pH 8.0, 1 mM EDTA, 50 mM NaCl), boiled in water and allowed to cool overnight at 4°C. Annealing was verified by running on a 4% agarose gel next to single stranded oligos. We generated ^32^P-labeled probes with T4 polynucleotide kinase (PNK) and γ-^32^P ATP (30 pmol probe, 1X T4 PNK reaction buffer, 16.5 pmol γ-^32^P ATP, 20 units T4 PNK). Reactions were incubated at 37°C for 30 min followed by heat inactivation at 65°C for 20 min. Unincorporated γ-^32^P ATP was removed by running the product through Illustra MicroSpin™ G-25 Columns (GE Healthcare) and probe radioactivity was measured.

All EMSA reactions consisted of binding buffer (10 mM Tris, 50 mM KCl, 1 mM DTT; pH 7.5), 5 mM MgCl_2_, 0.05 ug/ul Poly dI•dC, and 100,000 dpm of ^32^P-labeled probe. Each probe containing the predicted Tbx2 binding site was run in the presence and absence of cichlid Tbx2 (15 ug of nuclear lysate). Unlabeled wild-type and mutant probes (≥400 molar excess) were used to test the specificity of the binding. Reactions were incubated for 60 min at room temperature and run on a 6% DNA Retardation gel (Invitrogen™) in TBE buffer. Gels were then dried and exposed to radiographic film on an amplifying screen cassette overnight at −80°C.

### Luciferase Assays

To determine if Tbx2a directly affects expression of sequence downstream of the RH2 and LWS LCRs, we generated luciferase constructs. Because the RH2 LCR interacts with the promoter of all three downstream *RH2* genes (66) we chose to clone 495 bp of the RH2 LCR into pGL4.26, a luciferase plasmid that contains its own minimal promoter. For the LWS LCR we cloned the entire 3.9 kb region from the LWS LCR to the *LWS* transcriptional start site into pGL4.10; a promoter-less luciferase plasmid. For both plasmids; PCR products were generated from genomic DNA with a Phusion^®^ High-Fidelity PCR kit (NEB) and primers that introduced *KpnI* and *XhoI* restriction sites. Cloning of PCR products was performed as described above. For luciferase assays, 15,000 MDCK cells were plated into each well of a 24-well plate in 500μL of MEM alpha medium with 10% FBS. Two days after plating all wells were transfected with 150 ng of the respective reporter vector (empty pGL4.26, pGL4.26-RH2_LCR, empty pGL4.10, or pGL4.10-LWS), and 5 ng of thymidine kinase promoter-Renilla luciferase reporter plasmid (pRL-TK) as a control for transfection efficiency. Wells were co-transfected with either 0, 100, 250, 500, 750, or 1000ng of the pcDNA4-*Tbx2a* plasmid described above. Three replicate wells were run in each experiment and each experiment was repeated three times. An inert pUC19 plasmid was used to achieve transfections of equimolar amounts per well. Two days after transfection, cells were harvested, rinsed in PBS, lysed, and luciferase/*Renilla* signal was quantified on a GloMax®-Multi+ dual injector luminometer using a Dual-Glo® Luciferase Assay system (Promega). To determine the effect of increasing the amount of *Tbx2a* co-transfected on luciferase expression we transformed the measure of luciferase/*Renilla* for each co-transfected reaction relative to the average measure of luciferase/*Renilla* of the same plasmid that was not co-transfected with *Tbx2a*. To determine if Tbx2a was interacting with the RH2 LCR or *LWS* promoter we compared pGL4.26-RH2_LCR to empty pGL4.26 and pGL4.10-LWS to empty pGL4.10 at each treatment using t-tests in Prism v8.0.1.

## ACKNOWLEDGEMENTS

All animal procedures were approved by the University of Maryland College Park IACUC (Protocol # R-15-54). This work was supported by the National Eye Institute of the National Institutes of Health (R01EY024639 to K.L.C.; Intramural Research program ZIAEY000450 and ZIAEY000546 to A.S.).

## AUTHOR CONTRIBUTIONS

B.A.S., K.L.C., L.C., and A.S. designed the research; C.O. and B.A.S. generated the eQTL map and qPCR data; B.A.S. and S.P.N. generated sequence data; B.A.S. and L.C. performed protein experiments; B.A.S., M.C. and W.G. performed genome and transcriptome analyses. B.A.S. and K.L.C. wrote the manuscript; all authors revised the manuscript.

## DATA ACCESSIBILITY

DNA-seq and RNA-seq reads will be deposited at the NCBI Sequencing Read Archive. Other data has been included as supplementary materials.

## References

1. Cronin W, Johnsen S, Marshall J, Warrant J. Visual Ecology. Princeton: Princeton University Press; 2014.

2. Seehausen O, Terai Y, Magalhaes IS, Carleton KL, Mrosso HD, Miyagi R, et al. Speciation through sensory drive in cichlid fish. Nature. 2008;455(7213):620–6.

3. Sharpe LT, Gegenfurtner KR. Color Vision: From Genes to Perception. Cambridge, UK: Cambridge University Press; 1999.

4. Golding GB, Dean AM. The structural basis of molecular adaptation. Mol Biol Evol. 1998;15(4):355–69.

5. Yokoyama S, Yokoyama R. Adaptive Evolution of Photoreceptors and Visual Pigments in Vertebrates. Annu Rev Ecol Syst. 1996;27:543–67.

6. Yokoyama S. Molecular evolution of vertebrate visual pigments. Prog Retin Eye Res. 2000;19(4):385–419.

7. Carleton KL, Kocher TD. Cone opsin genes of african cichlid fishes: tuning spectral sensitivity by differential gene expression. Mol Biol Evol. 2001;18(8):1540–50.

8. Sandkam B, Young CM, Breden F. Beauty in the eyes of the beholders: colour vision is tuned to mate preference in the Trinidadian guppy (Poecilia reticulata). Mol Ecol. 2015;24(3):596–609.

9. Sandkam BA, Young CM, Breden FM, Bourne GR, Breden F. Color vision varies more among populations than among species of live-bearing fish from South America. BMC Evol Biol. 2015;15:225.

10. Fuller RC, Carleton KL, Fadool JM, Spady TC, Travis J. Population variation in opsin expression in the bluefin killifish, Lucania goodei: a real-time PCR study. J Comp Physiol A Neuroethol Sens Neural Behav Physiol. 2004;190(2):147–54.

11. Shand J, Davies WL, Thomas N, Balmer L, Cowing JA, Pointer M, et al. The influence of ontogeny and light environment on the expression of visual pigment opsins in the retina of the black bream, Acanthopagrus butcheri. J Exp Biol. 2008;211(Pt 9):1495–503.

12. Carleton KL, Spady TC, Streelman JT, Kidd MR, McFarland WN, Loew ER. Visual sensitivities tuned by heterochronic shifts in opsin gene expression. BMC Biol. 2008;6:22.

13. Fuller RC, Claricoates KM. Rapid light-induced shifts in opsin expression: finding new opsins, discerning mechanisms of change, and implications for visual sensitivity. Mol Ecol. 2011;20(16):3321–35.

14. Shand J, Hart NS, Thomas N, Partridge JC. Developmental changes in the cone visual pigments of black bream Acanthopagrus butcheri. J Exp Biol. 2002;205(Pt 23):3661-7.

15. Carleton KL, Dalton BE, Escobar-Camacho D, Nandamuri SP. Proximate and ultimate causes of variable visual sensitivities: Insights from cichlid fish radiations. Genesis. 2016;54(6):299–325.

16. O’Quin KE, Smith AR, Sharma A, Carleton KL. New evidence for the role of heterochrony in the repeated evolution of cichlid opsin expression. Evol Dev. 2011;13(2):193–203.

17. O’Quin KE, Schulte JE, Patel Z, Kahn N, Naseer Z, Wang H, et al. Evolution of cichlid vision via trans-regulatory divergence. BMC Evol Biol. 2012;12:251.

18. O’Quin KE, Smith D, Naseer Z, Schulte J, Engel SD, Loh YH, et al. Divergence in cisregulatory sequences surrounding the opsin gene arrays of African cichlid fishes. BMC Evol Biol. 2011;11:120.

19. Brawand D, Wagner CE, Li YI, Malinsky M, Keller I, Fan SH, et al. The genomic substrate for adaptive radiation in African cichlid fish. Nature. 2014;513(7518):375-+.

20. Behesti H, Papaioannou VE, Sowden JC. Loss of Tbx2 delays optic vesicle invagination leading to small optic cups. Dev Biol. 2009;333(2):360–72.

21. Takabatake Y, Takabatake T, Takeshima K. Conserved and divergent expression of T-box genes Tbx2-Tbx5 in Xenopus. Mech Dev. 2000;91(1-2):433-7.

22. Abrahams A, Parker MI, Prince S. The T-box transcription factor Tbx2: its role in development and possible implication in cancer. IUBMB Life. 2010;62(2):92–102.

23. Alvarez-Delfin K, Morris AC, Snelson CD, Gamse JT, Gupta T, Marlow FL, et al. Tbx2b is required for ultraviolet photoreceptor cell specification during zebrafish retinal development. Proceedings of the National Academy of Sciences. 2009;106(6):2023–8.

24. Tsujimura T, Chinen A, Kawamura S. Identification of a locus control region for quadruplicated green-sensitive opsin genes in zebrafish. P Natl Acad Sci USA. 1042007. p. 12813–8.

25. Tsujimura T, Hosoya T, Kawamura S. A single enhancer regulating the differential expression of duplicated red-sensitive opsin genes in zebrafish. PLoS Genet. 2010;6(12):e1001245.

26. Tam KJ, Watson CT, Massah S, Kolybaba AM, Breden F, Prefontaine GG, et al. Regulatory function of conserved sequences upstream of the long-wave sensitive opsin genes in teleost fishes. Vision Res. 2011;51(21-22):2295–303.

27. Hartl DL. Discovery of the transposable element Mariner. 2001;157(2):471–6.

28. Musser JM, Arendt D. Loss and gain of cone types in vertebrate ciliary photoreceptor evolution. Dev Biol. 2017;431(1):26–35.

29. Swaroop A, Kim D, Forrest D. Transcriptional regulation of photoreceptor development and homeostasis in the mammalian retina. Nat Rev Neurosci. 2010;11(8):563–76.

30. Liu F, Cao JX, Lv JH, Dong L, Pier E, Xu GX, et al. TBX2 expression is regulated by PAX3 in the melanocyte lineage. Pigm Cell Melanoma R. 2013;26(1):67–77.

31. Parry JW, Carleton KL, Spady T, Carboo A, Hunt DM, Bowmaker JK. Mix and match color vision: tuning spectral sensitivity by differential opsin gene expression in Lake Malawi cichlids. Curr Biol. 2005;15(19):1734–9.

32. Halfon MS. Perspectives on Gene Regulatory Network Evolution. Trends Genet. 2017;33(7):436–47.

33. Viets K, Eldred K, Johnston RJ, Jr. Mechanisms of Photoreceptor Patterning in Vertebrates and Invertebrates. Trends Genet. 2016;32(10):638–59.

34. Sebé-Pedrós A, Ruiz-Trillo I. Evolution and Classification of the T-Box Transcription Factor Family. In: Frasch M, editor. T-box Genes in Development and Disease. 122. Cambridge, MA: Academic Press; 2017. p. 1–26.

35. Sowden JC, Holt JK, Meins M, Smith HK, Bhattacharya SS. Expression of Drosophila omb-related T-box genes in the developing human and mouse neural retina. Invest Ophthalmol Vis Sci. 2001;42(13):3095–102.

36. Macosko EZ, Basu A, Satija R, Nemesh J, Shekhar K, Goldman M, et al. Highly Parallel Genome-wide Expression Profiling of Individual Cells Using Nanoliter Droplets. Cell. 2015;161(5):1202–14.

37. Li Q, Peterson KR, Fang X, Stamatoyannopoulos G. Locus control regions. Blood. 2002;100(9):3077–86.

38. Nandamuri SP, Conte MA, Carleton KL. Multiple trans QTL and one cis-regulatory deletion are associated with the differential expression of cone opsins in African cichlids. BMC Genomics. 2018;19.

39. Loosli F, Winkler S, Burgtorf C, Wurmbach E, Ansorge W, Henrich T, et al. Medaka eyeless is the key factor linking retinal determination and eye growth. Development. 2001;128(20):4035–44.

40. Munoz-Lopez M, Garcia-Perez JL. DNA transposons: nature and applications in genomics. Curr Genomics. 2010;11(2):115–28.

41. Conte MA, Joshi R, Moore EC, Nandamuri SP, Gammerdinger WJ, Roberts RB, et al. Chromosome-scale assemblies reveal the structural evolution of African cichlid genomes. Gigascience. 2019;8(4).

42. Conte MA, Gammerdinger WJ, Bartie KL, Penman DJ, Kocher TD. A high quality assembly of the Nile Tilapia (Oreochromis niloticus) genome reveals the structure of two sex determination regions. BMC Genomics. 2017;18(1):341.

43. Conte MA, Kocher TD. An improved genome reference for the African cichlid, Metriaclima zebra. BMC Genomics. 2015;16:724.

44. Untergasser A, Cutcutache I, Koressaar T, Ye J, Faircloth BC, Remm M, et al. Primer3-new capabilities and interfaces. Nucleic Acids Res. 2012;40(15).

45. Hofmann CM, O’Quin KE, Marshall NJ, Cronin TW, Seehausen O, Carleton KL. The eyes have it: regulatory and structural changes both underlie cichlid visual pigment diversity. PLoS Biol. 2009;7(12):e1000266.

46. Spady TC, Seehausen O, Loew ER, Jordan RC, Kocher TD, Carleton KL. Adaptive molecular evolution in the opsin genes of rapidly speciating cichlid species. Mol Biol Evol. 2005;22(6):1412–22.

47. Spady TC, Parry JW, Robinson PR, Hunt DM, Bowmaker JK, Carleton KL. Evolution of the cichlid visual palette through ontogenetic subfunctionalization of the opsin gene arrays. Mol Biol Evol. 2006;23(8):1538–47.

48. Dalton BE, Loew ER, Cronin TW, Carleton KL. Spectral tuning by opsin coexpression in retinal regions that view different parts of the visual field. Proc Biol Sci. 2014;281(1797).

49. Dalton BE, de Busserolles F, Marshall NJ, Carleton KL. Retinal specialization through spatially varying cell densities and opsin coexpression in cichlid fish. J Exp Biol. 2017;220(Pt 2):266–77.

50. Altschul SF, Madden TL, Schaffer AA, Zhang JH, Zhang Z, Miller W, et al. Gapped BLAST and PSI-BLAST: a new generation of protein database search programs. Nucleic Acids Res. 1997;25(17):3389–402.

51. Johnson M, Zaretskaya I, Raytselis Y, Merezhuk Y, McGinnis S, Madden TL. NCBI BLAST: a better web interface. Nucleic Acids Res. 2008;36:W5–W9.

52. Broman KW, Wu H, Sen S, Churchill GA. R/qtl: QTL mapping in experimental crosses. Bioinformatics. 2003;19(7):889–90.

53. Team RC. R: A language and environment for statistical computing. Vienna, Austria: R Foundation for Statistical Computing; 2018.

54. Wu TD, Watanabe CK. GMAP: a genomic mapping and alignment program for mRNA and EST sequences. Bioinformatics. 2005;21(9):1859–75.

55. Trapnell C, Roberts A, Goff L, Pertea G, Kim D, Kelley DR, et al. Differential gene and transcript expression analysis of RNA-seq experiments with TopHat and Cufflinks. Nature Protocols. 2012;7:562.

56. Schulte JE, O’Brien CS, Conte MA, O’Quin KE, Carleton KL. Interspecific variation in Rx1 expression controls opsin expression and causes visual system diversity in African cichlid fishes. Mol Biol Evol. 2014;31(9):2297–308.

57. Katoh K, Asimenos G, Toh H. Multiple alignment of DNA sequences with MAFFT. Methods Mol Biol. 2009;537:39–64.

58. Bazinet AL, Zwickl DJ, Cummings MP. A Gateway for Phylogenetic Analysis Powered by Grid Computing Featuring GARLI 2.0. Syst Biol. 2014;63(5):812–8.

59. Bolger AM, Lohse M, Usadel B. Trimmomatic: a flexible trimmer for Illumina sequence data. Bioinformatics. 2014;30(15):2114–20.

60. Li H, Durbin R. Fast and accurate long-read alignment with Burrows–Wheeler transform. Bioinformatics. 2010;26(5):589–95.

61. Robinson JT, Thorvaldsdottir H, Winckler W, Guttman M, Lander ES, Getz G, et al. Integrative genomics viewer. Nat Biotechnol. 2011;29(1):24–6.

62. Wittkopp PJ, Haerum BK, Clark AG. Evolutionary changes in cis and trans gene regulation. Nature. 2004;430:85.

63. Kohany O, Gentles AJ, Hankus L, Jurka JJBB. Annotation, submission and screening of repetitive elements in Repbase: RepbaseSubmitter and Censor. 2006;7(1):474.

64. Schwartz S, Zhang Z, Frazer KA, Smit A, Riemer C, Bouck J, et al. PipMaker—A Web Server for Aligning Two Genomic DNA Sequences. Genome Res. 2000;10(4):577–86.

65. Mathelier A, Fornes O, Arenillas DJ, Chen C-y, Denay G, Lee J, et al. JASPAR 2016: a major expansion and update of the open-access database of transcription factor binding profiles. Nucleic Acids Res. 2016;44(D1):D110–D5.

66. Tsujimura T, Masuda R, Ashino R, Kawamura SJBG. Spatially differentiated expression of quadruplicated green-sensitive RH2 opsin genes in zebrafish is determined by proximal regulatory regions and gene order to the locus control region. 2015;16(1):130.

