## Supplementary for "*Tbx2a* modulates switching of opsin gene expression"

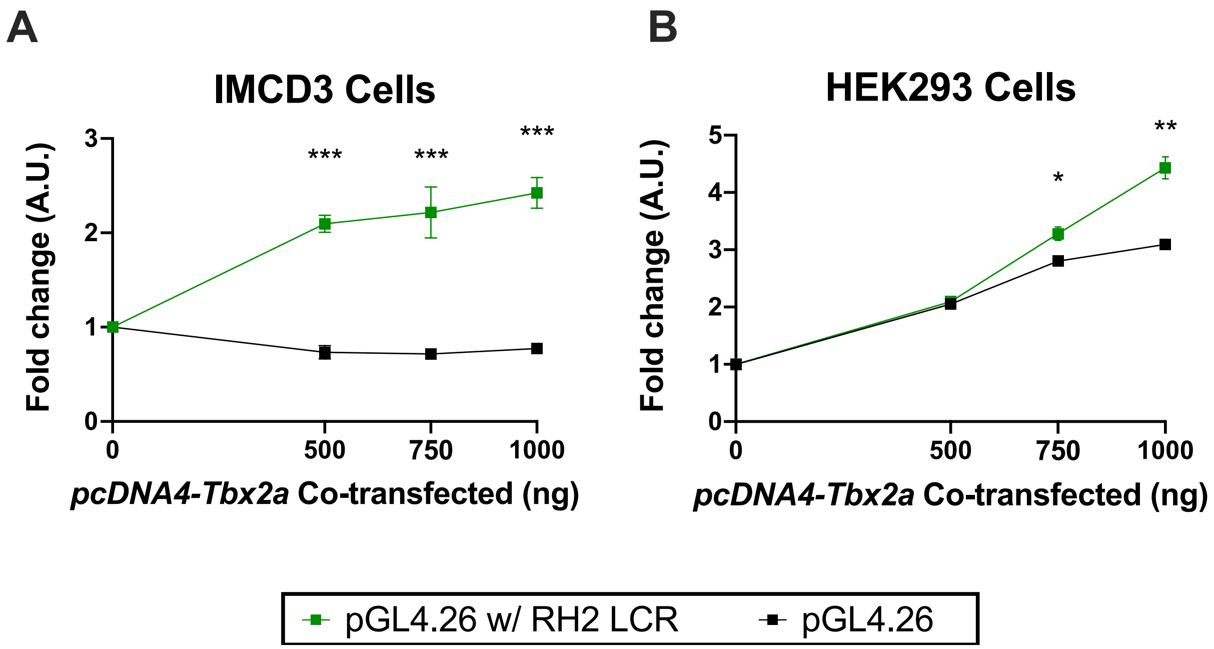


**Supplementary Figure 1.** Luciferase results of the Tbx2a interaction with the RH2 LCR were tested in both HEK293 and IMCD3 cells following the protocol used with MDCK cells with two minor deviations: (1) a pRL-TK renilla plasmid was used to calibrate luciferase signal rather than a pRL-CMV renilla plasmid, and (2) we did not conduct 100ng and 250ng co-transfections.

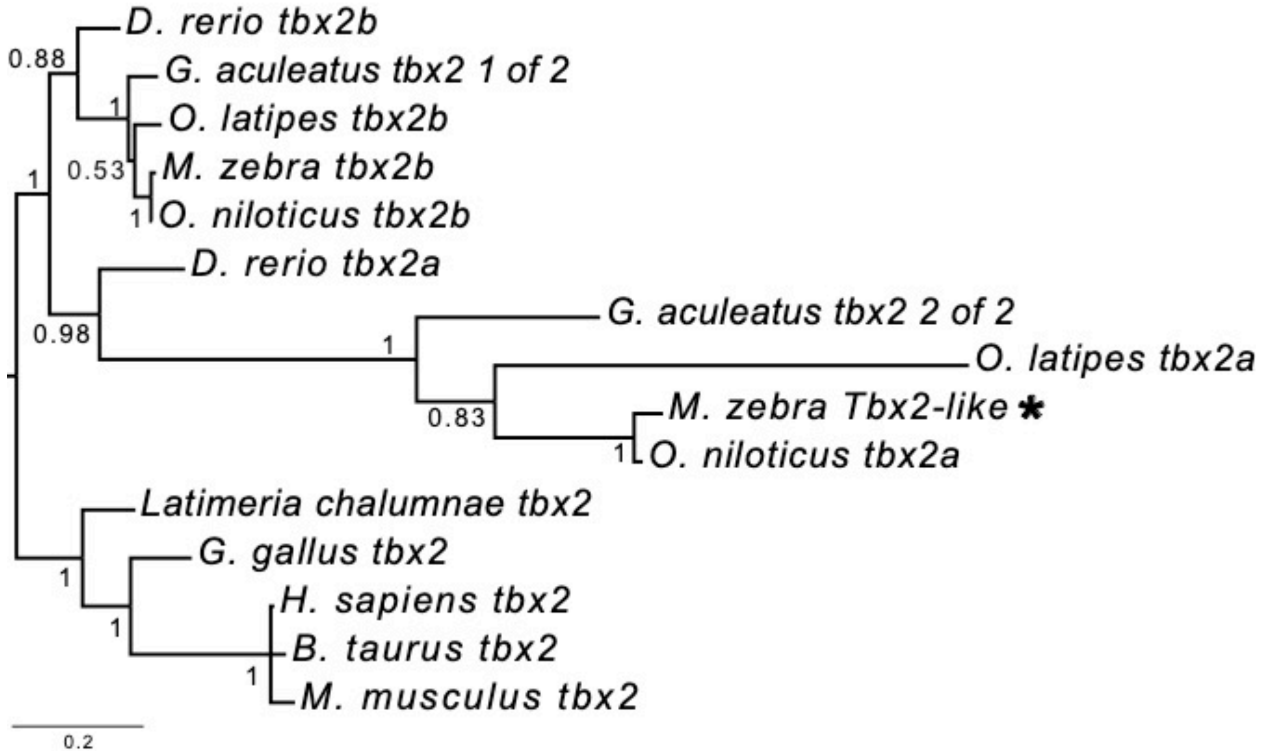


Supplementary Figure 2. Maximum likelihood tree built for *Tbx2* genes across eight species. The 'Tbx2-like' gene within the *LWS*/*RH2A* eQTL (starred) falls within the *Tbx2a* clade.

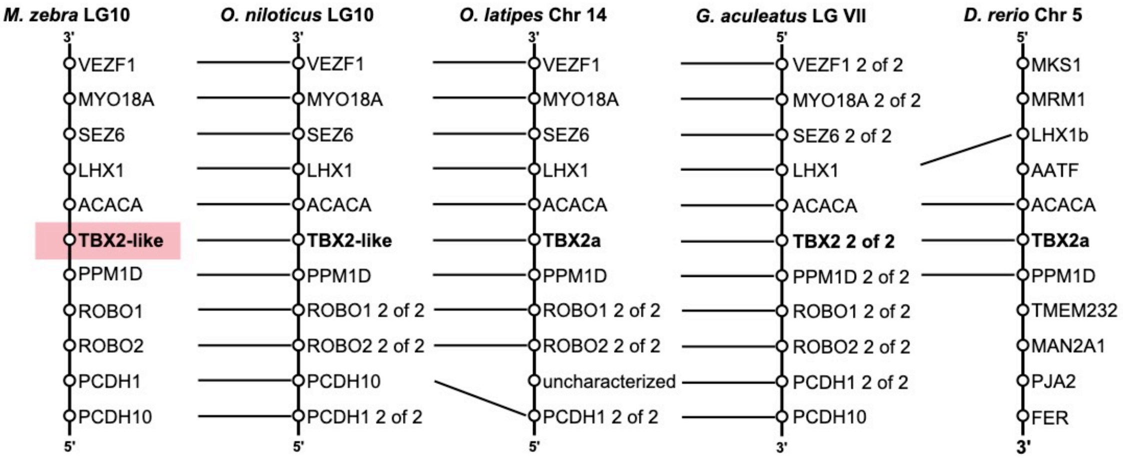


Supplementary Figure 3. Synteny of 'Tbx2-like' confirms the identity of 'Tbx2-like' to be *Tbx2a*.

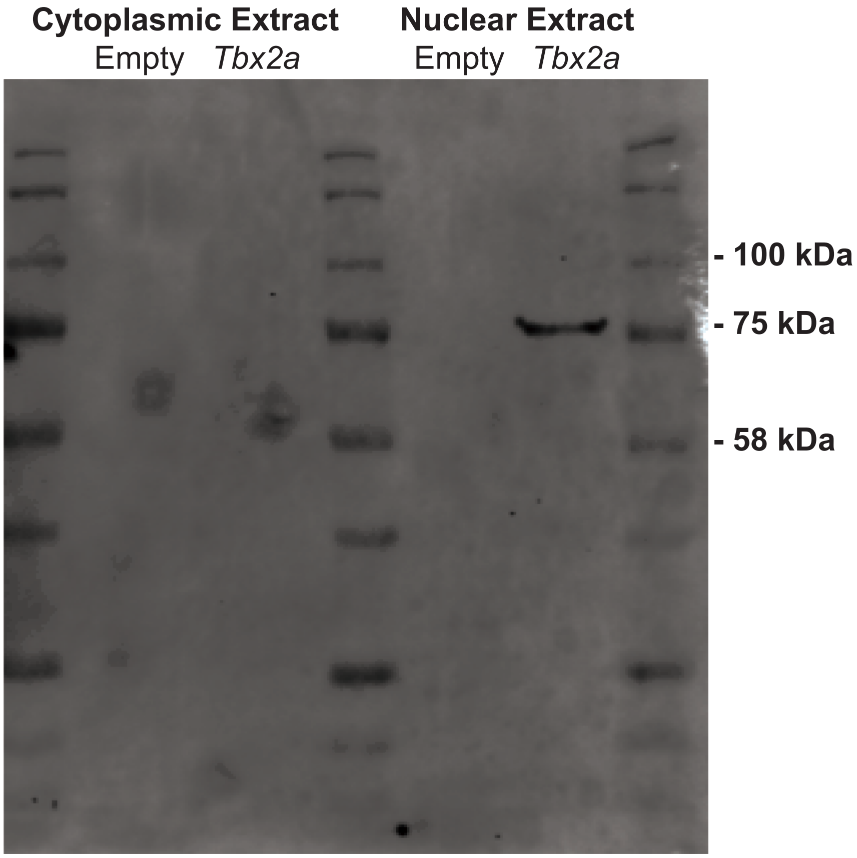


Supplementary Figure 4. Nuclear localization of Tbx2a in MDCK cells. Nuclear and cytoplasmic protein fractions were obtained from MDCK cells transfected with Tbx2 expression plasmids (+Tbx2a) or plasmids without the *Tbx2* coding region sequence (-Tbx2a) and subjected to Western blotting.

**Supplementary Table 1.** Differential expression between *T. intermedius* / *A. baenschi* and correlation with *LWS* across 5 cichlid species for each of the 31 genes within the *LWS*/*RH2A* eQTL. *NA* in the ratio of *T. intermedius* / *A. baenschi* expression indicates neither species expressed that gene. *NA* in the correlation across 5 cichlid species indicates less than two species expressed that gene.

| **NCBI accession** | **Gene Name** | **Expression ratio between *T. intermedius* and *A. baenschi* (F-1)^2** | **Correlation with *LWS* across 5 cichlid species** |
|---|---|---|---|
| XM_014408604.2 | ubiquitin C (*ubc*) | 0.300 | 0.918 |
| XM_004557411.5 | centromere protein V, transcript variant X2 (*cenpv*) | 0.011 | 0.741 |
| XM_004557412.2 | phosphatidylinositol glycan anchor biosynthesis class L (*pigl*) | 0.052 | 0.190 |
| XM_004557415.4 | nuclear receptor corepressor 1, transcript variant X3 (*ncor1*) | 0.042 | 0.130 |
| XM_004557409.4 | zinc finger SWIM-type containing 7 (*zswim7*) | 0.123 | 0.225 |
| XM_004557408.5 | adenosine A2b receptor (*adora2b*) | 5.035 | 0.012 |
| XR_002721188.2 | sperm antigen with calponin homology and coiled-coil domains 1, transcript variant X9 (*specc1*) | 0.127 | 0.022 |
| XM_004557407.5 | translocase of inner mitochondrial membrane 22 (*timm22*) | 0.008 | 0.000 |
| XM_004557402.2 | TBC1 domain family member 8B, transcript variant X2 (*tbc1d8b*) | 0.323 | 0.197 |
| XM_004557400.3 | gap junction alpha-3 protein-like (*gja3)* | *NA* | *NA* |
| XM_004557398.3 | gap junction beta-1 protein, transcript variant X1 (*gjb1*) | 2.127 | 0.085 |
| XM_004557397.3 | oligophrenin 1 (*ophn1*) | 0.156 | 0.059 |
| XM_004557395.5 | androgen receptor | *NA* | 0.190 |
| XM_024803896.1 | uncharacterized LOC112435397, transcript variant X2 | *NA* | 0.679 |
| XM_012920822.3 | moesin, transcript variant X2 (*msn*) | 0.380 | 0.022 |
| XM_004557393.3 | zinc finger CCCH-type containing 12B, transcript variant X3 (*zc3h12b*) | 0.756 | 0.147 |

| | | | |
|---|---|---|---|
| XM_004557390.3 | APC membrane recruitment protein 1 (*amer1*) | 0.157 | 0.000 |
| XM_004557388.4 | GRB2 associated binding protein 3, transcript variant X1 (*gab3*) | 0.596 | 0.043 |
| XM_004557387.4 | apoptosis inducing factor mitochondria associated 1 (*aifm1*) | 0.020 | 0.028 |
| XM_004557385.3 | glycine receptor subunit alpha-2 (*glra2*) | 0.143 | 0.147 |
| XM_004557386.2 | NUFIP2, FMR1 interacting protein 2 (*nufip2*) | 0.055 | 0.040 |
| XM_004557377.2 | musashi RNA binding protein 2, transcript variant X2 (*msi2*) | 1.863 | 0.000 |
| XM_004557373.2 | oligodendrocyte-myelin glycoprotein-like (*omgp*) | *NA* | 0.047 |
| XM_004557372.2 | uncharacterized LOC101484605 | 1.534 | 0.038 |
| XM_004557368.3 | kinase suppressor of ras 1, transcript variant X1 (*ksr1*) | 0.005 | 0.018 |
| XM_004557367.5 | vascular endothelial zinc finger 1 (*vezf1*) | 0.122 | 0.684 |
| XM_023152781.2 | unconventional myosin-XVIIIa, transcript variant X6 (*myo18a*) | 0.337 | 0.297 |
| XM_004557362.2 | seizure protein 6 homolog, transcript variant X2 (*sez6*) | 0.042 | 0.012 |
| XM_004557360.2 | LIM/homeobox protein Lhx1-like (*lhx1*) | *NA* | *NA* |
| XM_014408616.3 | acetyl-CoA carboxylase alpha, transcript variant X4 (*acaca*) | 0.048 | 0.344 |
| XM_004557464.1 | T-box transcription factor TBX2b-like (*Tbx2a*) | 105.954 | 0.878 |

**Supplementary Table 2.** Probe sequences and results of Electrophoretic Mobility Shift Assays (EMSA). Highlighted region denotes bases changed in the mutant probes.

| Probe | Sequence | Tbx2a binding in EMSA |
|---|---|---|
| RH2 LCR | TTACGTACAAGGTGTGAATCCAATTTGTAGGGATTGAGGATT | Yes |
| RH2 LCR Mutant | TTACGTACAAGAACGGAATCCAATTTGTAGGGATTGAGGATT | No |
| RH2Ab Promoter | ACCTGTGTGTGGTTGGTTTACACCTTTCTAACACTACTGGG | No |
| RH2Ab Promoter Mutant | ACCTGTGTGTGGTTGGTTTCGTTCTTTCTAACACTACTGGG | No |
| LWS LCR 1 | CTTCGCTCACTTTTCACTGTGTCAAATAGGTTAGACATTCA | No |
| LWS LCR 1 Mutant | CTTCGCTCACTTTTCACTAACGCAAATAGGTTAGACATTCA | No |
| LWS LCR 2 | TGTTCCGGAGAGATTAAAGTGTTTACGGAGCGAGCATGCCA | No |
| LWS LCR 2 Mutant | TGTTCCGGAGAGATTAAAAACGTTACGGAGCGAGCATGCCA | No |
| LWS Promoter 1 | ACTTTTGAATACAGAGAGGTGTCAAATCCAGACACTTTGCA | Yes |
| LWS Promoter 1 Mutant | ACTTTTGAATACAGAGAAACCCTGAATCCAGACACTTTGCA | No |
| LWS Promoter 2 | GCATGAGTTATTGTTAACGTGTGAAGTTGGATTTCATAAAC | No |
| LWS Promoter 2 Mutant | GCATGAGTTATTGTTAACAACGGAAGTTGGATTTCATAAAC | No |

Supplementary Table 3. Fixed differences between *T. intermedius* and *A. baenschi* between *Tbx2a* and the next gene upstream; Protein Phophatase 1D (~19.5kb), and the next gene downstream; Acetyl-CoA carboxylase alpha (~5.5kb). The 20 bp before and 20 bp after the fixed variant have been added to each sequence. Yellow denotes base pairs that differ between *T. intermedius* and *A. baenschi*. Blue denotes sequence that is present in only one species.

| Variant | Sequence + 20 bp up & 20 bp down |
|---|---|
| Upstream_1_*A_baenschi* | AAATGCAGTTGGCGCCATTGCTCCCATGGCTTATTATCCTT |
| Upstream_1_*T_intermedius* | AAATGCAGTTGGCGCCATTGTTCCCATGGCTTATTATCCTT |
| Upstream_2_*A_baenschi* | GAAGGTCCTGGATTTGAATACACCATCTGGCGCTTCTTGTG |
| Upstream_2_*T_intermedius* | GAAGGTCCTGGATTTGAATATACCATCTGGCGCTTCTTGTG |
| Upstream_3_*A_baenschi* | AAGCCCCCTAAATGAATTTGTAGGAAAGCAGAGCTTACTGC |
| Upstream_3_*T_intermedius* | AAGCCCCCTAAATGAATTTGAAGGAAAGCAGAGCTTACTGC |
| Upstream_4_*A_baenschi* | TTCCACAGCATGTCTTCTATGCTGTCTTCCTTCTCTCACCC |
| Upstream_4_*T_intermedius* | TTCCACAGCATGTCTTCTATCCTGTCTTCCTTCTCTCACCC |
| Upstream_5_*A_baenschi* | AGTCAGAATTTTCTCGTCACTCACCATGGGGTGAATCCAGT |
| Upstream_5_*T_intermedius* | AGTCAGAATTTTCTCGTCACCCACCATGGGGTGAATCCAGT |
| Upstream_6_*A_baenschi* | TCACCCATAGGCGCGCGCGCACACACACACACACACACACA |
| Upstream_6_*T_intermedius* | TCACCCATAGGCGCGCGCGCGCACACACACACACACACACA |
| Upstream_7_*A_baenschi* | CCCACTTAGTCACCTGACATCTGATCGACCCATCCCCAAAT |
| Upstream_7_*T_intermedius* | CCCACTTAGTCACCTGACATTTGATCGACCCATCCCCAAAT |
| Upstream_8_*A_baenschi* | CAGCTCGTCCCGGTCCAGCCCGCCTCCCGCTCGCCTCTGCC |
| Upstream_8_*T_intermedius* | CAGCTCGTCCCGGTCCAGCCTGCCTCCCGCTCGCCTCTGCC |
| Upstream_9_*A_baenschi* | AGGCTCACCATGAGTGATAGGTGGATGCTCAAAACCAGTTA |
| Upstream_9_*T_intermedius* | AGGCTCACCATGAGTGATAGTTGGATGCTCAAAACCAGTTA |
| Upstream_10_*A_baenschi* | AAAAATGGCTAAATTAAATCAATAAGGTCGGGTTTAGTAACGTATCG CTGT |

| | |
|---|---|
| Upstream_10_*T_intermedius* | AAAAATGGCTAAATTAAATCAATTTAGCAAATAAGAGTGGGCCTAAA<br>TTCCTCGACACTGAGGTGAATGACTCACTTCCAGGTGTTGCAAAGGC<br>TTGATTGCAGTTCTGGCTGCCAAGGGTGGCACAACCACTTATTAACT<br>TTAGGTGAAATTACTTTTTCACATAGGGCCAGGTAGGTTTAATAAAA<br>GAAATCATCATTCTAAAACTGTATTTTCTATTTATTCAAGTTACTTT<br>CTCTAATATTAAAATTTGTTGGTGATATGATGCAAGTAAGTGTGGCA<br>AATATGCAAAGAACCAAGAAATTTGGGGTGTGGGGTAAATACTTTTT<br>TACAGCACTATACGAACTCACGTGGCTTACTCACTGTATCCTCAAAC<br>CCTCCAGCCATATTCTGTTTTCCAGGAATAGGTGAACTGCTTGAGAA<br>AGGTTTGCCTGTTGATCACAAAATCCATTTAACTCTGAGCCAGGTGT<br>CAATGTGCCCAAGTGTACAATGAATAATGCTCCATCACATTATGAGC<br>AGACAGGCTTTGGCTAAAGTCAGCGAGACAGCATACTACACCAAATT<br>CCGATGGATAGAAATACTTTGACATATCATTACTATGATATGCTTTA<br>TGTTTTAAATATAAAATGTTAGTTTAAAAAAAGAAATACATTTAGAA<br>AAAAACTATTTATGCATTAATTACTTCATTTCTCAAGACAGGCATTC<br>TGACTTGGGAAACACATGTAAATAATAACACTTATTATATTCCATTT<br>TCCTGCTGTGATCTGTCACAGTGTCTAGACTGTTTGTTGAACTTGCC<br>GCAACAGACAATTTGAAATTACAGCCAGCTTGGGAGTAAGTGATGTG<br>ACCCAAGATTGGAGCAGGATTTAGAGGTATGGTGAGGTTAAGATTCT<br>TAACTGTCTCTAAGTCCTTTGATTGTTTCCCTGTAAAATCTTTTTCA<br>CCTCACACACATTACGAACAGTTTGAAGGCAGATTGACCTGCCCAGG<br>TATAAGGTCGGGTTTAGTAACGTATCGCTGT |
| Downstream_1_*A_baenschi*<br>Downstream_1_*T_intermedius* | CTCAACCGTGTGAAAAAAAATTTAGCTGGACAGTGACACTA<br>CTCAACCGTGTGAAAAAAAAATTAGCTGGACAGTGACACTA |
| Downstream_2_*A_baenschi*<br>Downstream_2_*T_intermedius* | CAGCCTTAGTTCTTGTTTTTTCCCCCCCTTAATTTTTCTTT<br>CAGCCTTAGTTCTTGTTTTYCCCCCCCCTTAATTTTTCTTT |
| Downstream_3_*A_baenschi*<br>Downstream_3_*T_intermedius* | CAGTTGCTCTGACAGGTGGGTTTTTTTTTAACAGTTGTGG<br>CAGTTGCTCTGACAGGTGGGTTTTTTTTTTAACAGTTGTGG |

Supplementary Table 4. qTR-PCR Assay primer/probes and efficiencies

| Gene | Efficiency | Forward Primer | Probe | Reverse Primer |
|---|---|---|---|---|
| *Tbx2a* | 0.8703 | TTGGGTTTAAGTCTGAGAAGTGAAGA | ACTGTGGAACAACATCACCTCTTCAGCTCC | TGCTGTGACAGTTGACGTTTGA |
| *Gnat2* | 0.8511 | AACGATGGTGCTCTTCCCC | GACTCACCAGCACCAAGCAGTAATAGCTTGA | CCAAGAAGATGCTGATAAGGAGTCT |
| *LWS* | 0.8541 | CTGTGCTACCTTGCTGTGTGG | TGGCCATCCGTGCTGTTGCC | GCCTTCTGGGTTGACTCTGACT |

Supplementary Table 5. Retinal qRT-PCR of wild caught Malawi species used for correlation between *Tbx2a* and *LWS* opsin gene expression.

| **Genus** | **Species** | ***Tbx2a / Gnat2*** | ***LWS / Gnat2*** |
|---|---|---|---|
| *Tyrranochromis* | *macrostoma* | 0.004552 | 5.817233 |
| *Copadichromis* | *eucinostomus* | 0.008268 | 2.483387 |
| *Protomelas* | *taeniolatus* | 0.005806 | 5.156007 |
| *Petrotilapia* | *nigra* | 0.008992 | 2.276384 |
| *Stigmatochromis* | *woodi* | 0.007842 | 1.397935 |
| *Aristochromis* | *christyi* | 0.001373 | 0.039104 |
| *Pseudotropheus* | *microstoma* | 0.004581 | 0.937664 |
| *Pseudotropheus* | *tropheops red cheek* | 0.005581 | 0.177687 |
| *Metriaclima* | *aurora* | 0.003645 | 0.100585 |
| *Nimbochromis* | *polystigma* | 0.004771 | 0.21695 |
| *Metriaclima* | *callainos* | 0.002176 | 0.042951 |
| *Nimbochromis* | *linni* | 0.003661 | 0.016758 |
| *Tyrranochromis* | *maculiceps* | 0.006847 | 5.382013 |
| *Labeotropheus* | *fuelleborni* | 0.005222 | 0.27066 |
| *Aulonocara* | *hansbaenschi* | 0.006828 | 1.027899 |
| *Genyochromis* | *mento* | 0.003292 | 0.045738 |
| *Hemitilapia* | *oxyrhynchus* | 0.002896 | 0.089443 |
| *Labidochromis* | *blue sp* | 0.007436 | 2.811899 |
| *Labeotropheus* | *trewavasae* | 0.002021 | 0.056938 |
| *Melanochromis* | *auratus* | 0.001653 | 0.018512 |
| *Metriaclima* | *zebra* | 0.00333 | 0.529718 |
| *Melanochromis* | *B&Wjohanni* | 0.003678 | 0.254426 |
| *Cynotilapia* | *afra* | 0.002963 | 0.006142 |
| *Cyrtocara* | *moorii* | 0.03045 | 7.513729 |
| *Rhamphochromis* | *esox* | 0.005321 | 0.259276 |
| *Protomelas* | *annectens* | 0.003493 | 0.10317 |
| *Taeniolatus* | *preorbitalis* | 0.078262 | 65.75685 |
| *Trematocranus* | *placodon* | 0.008209 | 0.457386 |
| *Maravachromis* | *mola* | 0.012772 | 6.02061 |
| *Placidochromis* | *johnstoni* | 0.004251 | 0.297976 |
| *Tropheus* | *intermediate* | 0.004334 | 0.183576 |
| *Dimidiochromis* | *compressiceps* | 0.009336 | 6.6337 |
| *Cyathochromis* | *obliquidens* | 0.008181 | 3.485875 |

| | | | |
|---|---|---|---|
| *Aulonocara* | *sp.* | 0.007187 | 0.314865 |
| *Metriaclima* | *sp.* | 0.005754 | 0.650807 |
| *Metriaclima* | *livingstonii* | 0.003637 | 0.290745 |
| *Tropheops* | *orange chest* | 0.025936 | 6.855971 |
| *Protomelas* | *fenestratus* | 0.007768 | 0.440525 |
| *Pseudotropheus* | *heteropictus* | 0.003542 | 0.108031 |
| *Protomelas* | *similis* | 0.029292 | 18.877 |
| *Protomelas* | *spinolotus* | 0.008701 | 5.071511 |
| *Lethrinops* | *aurita* | 0.00554 | 7.972951 |
| *Lethrinops* | *placodon* | 0.019416 | 14.29966 |
| *Otopharynx* | *pictus* | 0.017163 | 3.161022 |
| *Oreochromis* | *sp.* | 0.02591 | 3.282212 |
| *Tropheops* | *gracilior* | 0.006655 | 0.657307 |
| *Tropheops* | *"broad mouth"* | 0.004506 | 0.166757 |
| *Labidochromis* | *gigas* | 0.003617 | 0.124624 |
| *Placidochromis* | *milomi* | 0.008499 | 6.62759 |
| *Otopharynx* | *heterodon* | 0.011036 | 4.263379 |
| *Tramitichromis* | *brevis* | 0.013971 | 6.413635 |
| *Dimidiochromis* | *kwinge* | 0.036795 | 34.0777 |
| *Rhamphochromis* | *sp.* | 0.01306 | 3.145881 |
| *Melanochromis* | *vermivorus* | 0.00444 | 0.192097 |
| *Melenochromis* | *parallelus* | 0.004377 | 0.182764 |
| *Lethrinops* | *auritus* | 0.004175 | 2.696787 |
| *Copadichromis* | *jacksoni* | 0.002999 | 0.210175 |
